# A robust two-sample Mendelian Randomization method integrating GWAS with multi-tissue eQTL summary statistics

**DOI:** 10.1101/2020.06.04.135541

**Authors:** Kevin J. Gleason, Fan Yang, Lin S. Chen

## Abstract

In the post-genome-wide association era, two-sample Mendelian Randomization (MR) methods have been applied to detect genetically-regulated risk factors for complex diseases. Two-sample MR considers single nucleotide polymorphisms (SNPs) associated with a putative exposure as instrumental variables (IVs) to assess the effect of the exposure on an outcome by leveraging two sets of summary statistics: IV-to-exposure and IV-to-outcome statistics from existing GWASs. Traditional MR methods impose strong assumptions on the validity of IVs, and recent literature has relaxed the assumptions allowing some IVs to be invalid but generally requiring a large number of nearly independent IVs. When treating expression-quantitative-trait-loci (eQTLs) as IVs to detect gene expression levels affecting diseases, existing methods are limited in applicability since the numbers of independent eQTLs for most genes in the genome are limited. To address those challenges, we propose a robust two-sample MR framework that requires fewer IVs and allows moderate IV correlations and some IVs to be invalid. This is achieved by leveraging existing multi-tissue eQTL summary statistics (multiple sets of IV-to-exposure statistics) and GWAS statistics in a mixed model framework. We conducted simulation studies to evaluate the performance of the proposed method and apply it to detect putative causal genes for schizophrenia.

## Introduction

For more than a decade, genome-wide association studies (GWAS) have uncovered tens of thousands of unique associations between single nucleotide polymorphisms (SNPs) and complex diseases/traits [1]. In the post-GWAS era, the next major challenge is to further understand the biological mechanisms underlying the observed associations and identify clinically actionable risk factors for various complex diseases/traits. Most of the disease/trait-associated SNPs have small effect sizes and reside in non-coding regions with unknown functions [2; 3]. In order to elucidate their mechanisms and functions, many efforts have been made to integrate GWAS summary statistics with other information (e.g., eQTL statistics) and to identify genetically-regulated risk factors (e.g., gene expression levels) for complex diseases. Those methods include transcriptome-wide association studies (TWAS) [4; 5; 6; 7], colocalization analyses [8; 9; 10; 11], two-sample Mendelian Randomization (MR) analysis [12; 13; 14; 15] and others.

Compared to other integrative genomic analyses, MR analysis has its unique advantages. It steps beyond association towards causation, aiming to identify modifiable risk factors (exposures) for complex diseases while allowing unmeasured confounders affecting both exposures and disease outcomes of interest. Specifically, MR methods consider SNPs with known associations with an exposure of interest as instrumental variables (IVs) [16; 17; 18; 19]. Since SNP genotypes were ‘Mendelian Randomized’ from parents to offspring during meiosis, they are assumed to be generally unrelated to external confounders. Under certain assumptions, SNPs can be used as IVs to estimate and test for the causal effects of an exposure on a disease outcome from observational data. Two-sample MR methods refer to the MR methods requiring only two sets of summary statistics, IV-to-exposure and IV-to-outcome association statistics from two independent sets of samples, and thus are widely used to recapitalize on existing summary statistics.

Traditional MR methods imposed strong assumptions on the validity of IVs [20]. A valid IV is a genetic variant that affects the complex disease through only the exposure of interest (no direct effect) and is independent of unmeasured confounders of the exposure and the disease outcome [21]. That is, there is no ‘horizontal pleiotropy’ [16] (a phenomenon where a genetic variant also affects the complex trait via other pathways not through the exposure) nor ‘correlated pleiotropy’ [22] (a phenomenon where a genetic variant affects both exposure and outcome through a heritable shared factor, i.e. IVs are associated with the confounder). See Figure 1 for an illustration. Note that valid IVs do not have to be the causal SNPs. Due to the pervasive pleiotropic effects of SNPs and linkage disequilibrium (LD) among SNPs in a region, it is commonly observed that SNPs may be associated with multiple molecular, intermediate and/or complex traits [23; 24; 25]. Both horizontal and correlated pleiotropy effects are prevalent in the genome. The inclusion of invalid IVs in traditional MR analyses may lead to biased causal effect estimation and inference. More recently, robust MR methods have been proposed to relax the assumptions by considering multiple IVs and allowing some to be invalid. Some methods allow up to half of the IVs being invalid but require individual-level genotype and phenotype data, which may limit the applicability of the methods [26]. Some methods require IVs to be nearly independent [13; 14; 27; 15] and/or require the number of IVs to be large [22; 28]. Those methods have been successfully applied to detect intermediate non-omics traits as exposures for complex diseases. For example, in detecting the protective effect of high-density lipoprotein cholesterol (HDL-C) on peripheral vascular disease, the suspected modifiable exposure HDL-C has many established GWAS SNPs as potential IVs [28].

**Figure 1:**
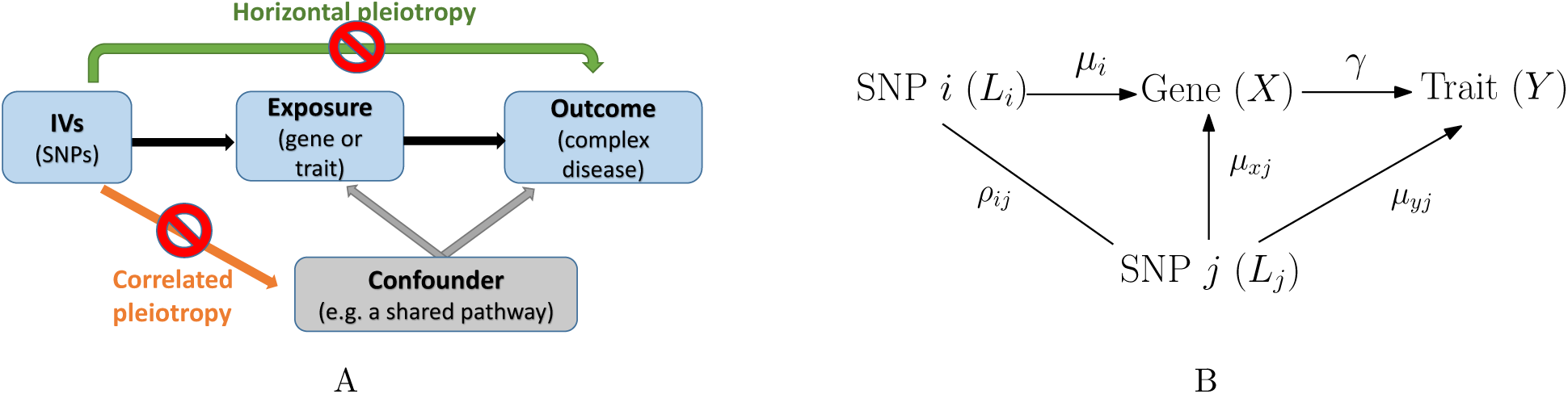
Illustrations of Mendelian Randomization analysis and assumptions. (A) When a SNP (or is in LD with a SNP that) is affecting the outcome not via the exposure of interest or is correlated with an unmeasured confounder for both the exposure and the outcome, the SNP is an invalid instrument. Note that the presence of unmeasured confounders is allowed in MR analysis, but instruments are assumed to be independent of the confounders. (B) An illustration of pleiotropy of SNP *j* in an LD block affecting the validity of SNP *i* of interest as an IV. A SNP *j* is in LD with an IV SNP *i* of interest. SNP *j* is an eQTL of the targeted gene and has a direct effect on the trait (horizontal pleiotropy). When conducting MR analysis with only marginal summary statistics, the effect of SNP *j* is not accounted for and will confound the relationships among the SNP *i*, the gene expression and the trait. That is, horizontal (and/or correlated) pleiotropy in a gene region will bias the effect estimate based on marginal statistics for SNP *i*, without conditioning on SNP *j*.

When applying MR methods to detect gene expression as an exposure for a disease outcome (termed as “transcriptome-wide MR” [29; 30]), new challenges arise. First, few studies have genotype, gene expression and disease outcome data being measured on the same set of samples, and even when all data is available for the same set of subjects, sample sizes are generally limited. Thus, MR methods requiring individual-level data may have limited power and applicability. Second, invalid IVs can be quite prevalent when studying gene expression as the exposure. Many genetic variants may affect complex diseases not completely via gene expression levels of a cis-gene [31]. Recent studies have reported the existence of many GWAS SNPs being also multi-omics QTLs (i.e., SNPs affecting both cis-gene expression and methylation levels then affecting complex diseases) [25; 24], and QTLs with effects on diseases mediated via splicing events [32]. Methods allowing invalid IVs are necessary in studying gene expression as the exposure. Last but foremost, when treating cis-eQTLs as IVs, the numbers of independent cis-eQTLs for most genes in the genome are very limited. Existing robust two-sample MR methods allowing invalid IVs generally require either multiple independent IVs or a large number of (weakly correlated) IVs, and those existing methods would have limited applicability in analyzing most genes in the genome.

To address those challenges in analyzing gene expression as the exposure for a disease outcome, we propose a two-sample Mendelian Randomization method ROBust to correlated and some INvalid instruments, termed “MR-Robin”. It requires only summary-level marginal GWAS and multi-tissue eQTL statistics as input, considers multi-tissue eQTL effects for multiple IVs of a gene, allows IVs to be correlated and some of them to be invalid, and can be applied to genes with only a small number of cis-eQTLs. Compared to existing two-sample MR methods allowing invalid IVs, MR-Robin lessens the required number of independent IVs by integrating GWAS statistics with multi-tissue eQTL statistics (i.e., multiple sets of IV-to-exposure summary statistics) in a mixed model framework. Moreover, by carefully selecting cross-tissue eQTLs as IVs, MR-Robin also improves the robustness of IV effects across “two-samples” and may improve the reproducibility of estimation and inference based on two-sample MR analyses. Specifically, MR-Robin considers the estimated effect of a gene on a disease from each IV as an observed value of the true effect plus a SNP-specific bias. By jointly considering multiple IVs, MR-Robin decomposes the estimated effects of multiple IVs into two components – a concordant effect shared across IVs and a discordant component allowing some IVs to be invalid with SNP-specific deviations from the true effect. MR-Robin makes the estimation identifiable by taking advantage of the multi-tissue eQTL effects for multiple IVs of a gene and treating them as the response variable in a reverse regression, with GWAS effect estimates as the predictor. The rich multi-tissue eQTL effect information in the response variable allows the estimation of SNP-specific random-slopes (i.e. deviated effects) due to potentially invalid IVs. Thus, with only a limited number of potentially correlated IVs, MR-Robin can test the effect from a gene to a disease by testing the shared (fixed effects) correlation between eQTL and GWAS effects across IVs. We conducted extensive simulations to evaluate the performance of MR-Robin under various scenarios in analyzing gene expression as the exposure for a disease outcome in the presence of invalid IVs. We applied MR-Robin to identify gene expression levels affecting schizophrenia risk by leveraging multi-tissue eQTL summary statistics from 13 brain tissues in the Genotype-Tissue Expression (GTEx) project [33] and GWAS summary statistics from the Psychiatric Genomics Consortium (PGC) [34].

## Methods

Let *β*_*xi*_ (*i* = 1, …, *I*) denote the marginal eQTL effect of a local eQTL/IV *i* for a gene and *β*_*yi*_ denote the marginal GWAS association effect on a complex trait of the eQTL/IV *i* in the GWAS study. Note that both *β*_*xi*_ and *β*_*yi*_’s are effects in the GWAS study, though *β*_*xi*_ is latent since expression data is not available for the GWAS samples, which is typical for most GWASs. Our goal is to test whether the effect of gene expression on the trait (*γ* in Figure 1) is zero, *H*_0_ : *γ* = 0 vs. *H*_*A*_ : *γ* ≠ 0. Traditional two-sample MR methods often take the ratio *β*_*yi*_*/β*_*xi*_ as an estimand for *γ* based on IV *i*. In the following subsections, we will first show that when there is a SNP *j* with a horizontal or correlated pleiotropic effect, and SNP *j* is in LD with the selected IV *i*, the ratio *β*_*yi*_*/β*_*xi*_ is a biased estimate for *γ* with a SNP-specific bias depending on many factors. Then we will introduce MR-Robin with a mixed model framework based on reverse regressions taking multi-tissue eQTL summary statistics (multiple sets of IV-to-exposure statistics) as response and IV-to-outcome statistic as predictor to test for a non-zero effect from gene to disease.

### Bias in *β*_*yi*_*/β*_*xi*_ as an estimand for *γ* when SNP with pleiotropy is in LD with IV *i*

Without loss of generality, we assume that there are two SNPs *i* and *j* in LD, and SNP *i* is a valid IV if conditioning on SNP *j*, and SNP *j* has a horizontal pleiotropic effect as depicted in Figure 1. For multiple eQTLs in an LD block, one can consider them as being conditionally valid IVs and invalid IVs. Below are the data generating models in a GWAS:

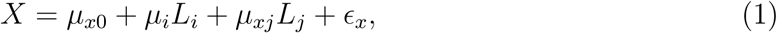

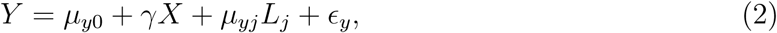

where *X* is the gene expression levels and *Y* is the continuous complex trait of interest in a GWAS study; and *L*_*i*_ and *L*_*j*_ are the genotypes for SNPs *i* and *j*, respectively. As a valid IV given *L*_*j*_, the genotype of SNP *i* (*L*_*i*_) is independent of the error terms *ϵ*_*x*_ and *ϵ*_*y*_. In the above models, the conditional association between *X* and *L*_*i*_ given *L*_*j*_ is captured by *µ*_*i*_, and the conditional association between *Y* and *L*_*i*_ given *L*_*j*_ is *γ · µ*_*i*_. And the ratio of the two, 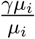, recovers the true effect of interest, *γ*.

Without adjusting for SNP *j*, the summary statistics for SNP *i* are calculated based on the following marginal models:

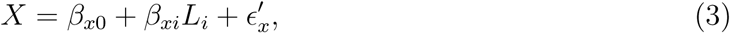

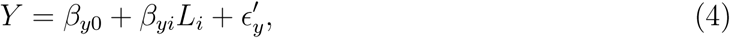

where *β*_*xi*_ and *β*_*yi*_ are the marginal eQTL and GWAS association effects, respectively, in the GWAS study. Note that one could also adjust covariates in the above models (1)-(4) and that does not affect our conclusion. We ignore covariates for simplicity. Define 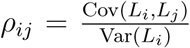, in terms of parameters in (1) and (2), it can be derived that the marginal effects 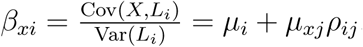, and 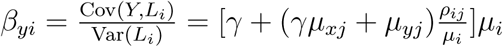.

It can be seen that the bias of marginal eQTL effect estimate for SNP *i* on gene expression, *β*_*xi*_, with respect to the true eQTL effect, *µ*_*i*_, is *µ*_*xj*_*ρ*_*ij*_. And the bias of marginal GWAS effect estimate for SNP *i* on complex trait, *β*_*yi*_, with respect to the mediated effect from SNP to gene to trait, *γµ*_*i*_, is (*γµ*_*xj*_ + *µ*_*yj*_)*ρ*_*ij*_. And it can be derived that the bias of the ratio of marginal GWAS to eQTL effect estimates, *β*_*yi*_*/β*_*xi*_, with respect to the true effect, *γ*, is given by 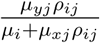. All the biases are functions of SNP *i*’s eQTL effect size, LD strength to the pleiotropic SNP *j* and effect size of the pleiotropy. Therefore, the bias will vary from SNP to SNP. Similarly, in the presence of correlated pleiotropic SNPs being in LD, the bias will also vary from SNP to SNP.

In the presence of horizontal or correlated pleiotropy in the LD region, an eQTL would be an invalid IV. And in such a case, the effect from gene to trait (*γ*) is not separable/identifiable from the direct effect of the eQTL nor confounding effects when only the total effect estimate (marginal summary statistic) is available. The presence of horizontal or correlated pleiotropy makes it challenging to infer the effect of a gene on a trait using single-IV-based MR approaches. When there are multiple eQTLs in the gene region, as shown in Figure 1, the presence of one SNP with horizontal or correlated pleiotropic effect would also render all eQTLs invalid if they are in LD.

It should be noted that the above bias is derived for analyzing gene expression as exposure for disease outcome based on marginal eQTL statistics. Due to the fact that all IVs (cis-eQTLs) are from the same cis-region and are in LD, the bias caused by pleiotropy in the region is particularly pronounced. When analyzing intermediate non-omics trait as the exposure and there are many known susceptibility loci from different genomic regions being associated with the non-omics exposure of interest, the IVs are generally less dependent and the bias due to local pleiotropy is generally specific to each locus.

### MR-Robin – a reverse-regression-based mixed model framework with multi-tissue eQTL statistics as response

Given the bias derived for *β*_*yi*_*/β*_*xi*_ w.r.t *γ*, we model that *β*_*yi*_*/β*_*xi*_ = (*γ* + *γ*_*i*_), where *γ*_*i*_ denotes the SNP-specific bias. The bias is zero if there is neither a horizontal nor correlated pleiotropic effect in the region. The bias is small to negligible for some eQTLs if those eQTLs themselves are valid IVs when adjusting for invalid IV *L*_*j*_, those eQTLs are in moderate-to-weak LD with the invalid IV(s), and the pleiotropic effect of SNP *j* is not strong (i.e., small *ρ*_*ij*_ *· µ*_*yj*_). It follows that

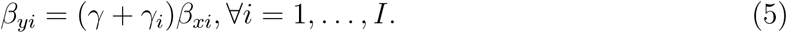

And equivalently,

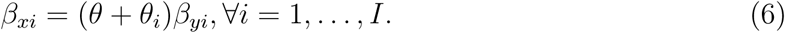

where *θ* captures the dependence between *β*_*xi*_ and *β*_*yi*_, and *θ*_*i*_ is the SNP-level deviation from the shared effect *θ* in the presence of pleiotropy.

In the above equation, *β*_*xi*_ is the marginal eQTL effect of SNP *i* to gene expression in the GWAS study and is often not available, since most GWAS studies do not have gene expression data measured. The availability of multi-tissue eQTL summary statistics from trait-relevant tissue types in a reference eQTL study such as GTEx provides a valuable resource to estimate *β*_*xi*_, given many cis-eQTL effects are shared across tissue types and are replicable across studies.

We model SNP *i*’s eQTL effect in tissue *k* (*k* = 1, …, *K*) in the reference multi-tissue eQTL data as a function of the eQTL effect in the GWAS data (*β*_*xi*_) and an error term. Based on (6), we propose the following model of MR-Robin for testing trait-association of a gene using only summary statistics from GWAS and multi-tissue eQTL reference:

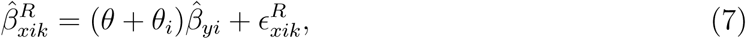

where 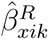 is the marginal eQTL effect estimate of the cross-tissue IV/eQTL *i* (*i* = 1, …, *I*) in the *k*-th tissue with the cross-tissue effect in the reference eQTL data, and 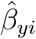 is the marginal GWAS effect estimate for SNP *i*; and *θ* captures the shared correlation of GWAS and eQTL statistics among all SNPs and is non-zero and bounded if and only if the true effect from the gene on the complex trait, *γ*, is non-zero and bounded; *θ*_*i*_ represents the SNP-specific bias due to horizontal or correlated pleiotropy in the region and is a SNP-specific random-slope; and 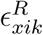 is a random error that follows a multivariate normal distribution 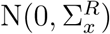. Note that there are both SNP-SNP correlations due to LD and tissue-tissue correlations due to sample overlapping. In the *P* -value estimation procedure, we account for the correlated errors by resampling.

In the reverse regression (7), the eQTL effect estimates from multiple tissue types, 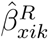, are considered as the response variable while the GWAS association effects 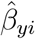 are considered as the predictor. This is mainly to take advantage of the rich information in multi-tissue eQTL datasets (i.e., variation in response). If there are multiple sets of correlated or independent GWAS summary statistics from the same population/ethnicity without study heterogeneity, often consortium-based meta-analysis may have been conducted with improved power and precision, and a single set of GWAS summary statistics would be made available. Each observation in the regression (7) is an estimated/observed marginal eQTL effect, with a total of *I* × *K* (SNP-by-tissue) observations. By testing the shared correlation of tissue-specific eQTL effects and the corresponding GWAS association effects for multiple eQTLs in the same gene (*H*_0_ : *θ* = 0 vs. *H*_*A*_ : *θ* ≠ 0) while also allowing for SNP-level deviation, we can test the effect of gene expression on trait (*H*_0_ : *γ* = 0 vs. *H*_*A*_ : *γ* ≠ 0), allowing invalid and correlated IVs.

Many existing methods in the MR literature allowing invalid IVs [22; 14; 26] include an intercept or a random intercept in the model to capture the direct effect from genotype to trait, i.e. horizontal pleiotropy. That is, SNP-to-disease association effects from GWAS are modeled as *β*_*yi*_ = *γ · β*_*xi*_ + *γ*_*i*_, ∀*i* = 1, …, *I*. The model fits better when individual-level data are available and statistics conditional on other SNPs in the region can be obtained or when summary statistics from joint models of multi-SNPs in the region are available. In contrast, in the MR-Robin model, there is no intercept nor random intercept. Instead, we include a random slope for each SNP to capture the effect due to potential pleiotropy in the region. This is because, by allowing correlated IVs and considering all eQTLs in a region, as shown above when there is a non-zero pleiotropic effect, most of the SNPs in the LD region would be affected with a non-zero (but possibly negligible) SNP-specific deviation *θ*_*i*_. Allowing correlated IVs and some invalid IVs even when the number of IVs are limited is also a major innovation of our model. Due to limited numbers of eQTLs/IVs for most genes in the genome, a model with both an intercept and a random slope may not be identifiable and thus is not explored.

To account for uncertainty in the eQTL effect estimation, we perform a weighted mixed-effects regression analysis and weight each “observation” (i.e., a tissue-specific eQTL effect) by the reciprocal of the estimated standard error for 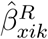, i.e., 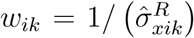. We obtain the *t*-statistic for testing the fixed effect of interest *θ* as our test statistic. To obtain the *P* -value while accounting for LD and tissue-tissue correlation as well as the uncertainty in the estimation of *β*_*yi*_’s, we adopt a resampling-based approach to generate the null test statistics. In each resampling *b* (*b* = 1, …, *B*), we sample a vector of GWAS effects from a multivariate distribution, 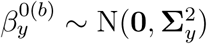, where the diagonal and off-diagonal elements are 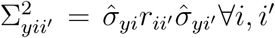 with *r*_*ii*′_ being the genotype correlation and 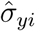 being the estimated standard error for 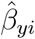. We apply the same weighted model (7) on data 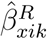 and 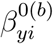’s to obtain a null statistic. We repeat the resampling process at least *B* = 10, 000 times and calculate the *P* -value. The MR-Robin algorithm is summarized in the algorithm below.

#### Algorithm 1

MR-Robin for assessing the causal effect of gene expression of a gene on a complex trait with summary statistics from GWAS and a multi-tissue eQTL study

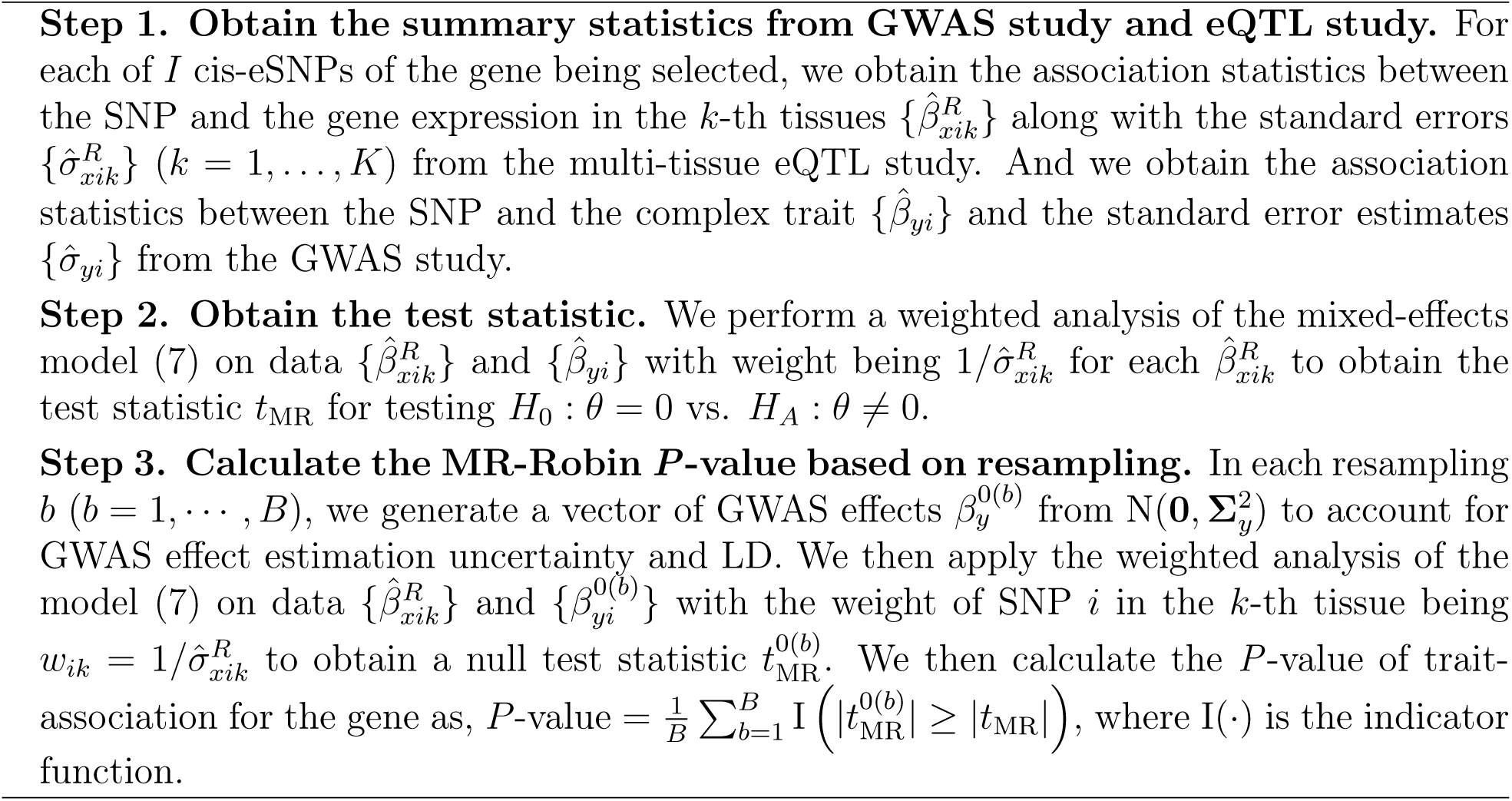

## Results

### Simulations to evaluate the performance of MR-Robin when IVs are correlated, some being invalid, and/or limited in number

In this section, we conducted simulation studies to evaluate the performance of MR-Robin as a two-sample MR method in the settings where a limited number of potentially correlated and/or invalid genetic variants are available as candidate instrumental variables (IVs). We showed that with multi-tissue eQTL statistics as input, MR-Robin is robust to the inclusion of correlated and some proportions of invalid IVs even when the number of IVs is small. We compared MR-Robin to several existing MR methods in the literature that are based on single-tissue eQTL and GWAS summary statistics and are robust to invalid IVs: MR-Egger [27], MR-RAPS [14], MRMix [13], and BWMR (a Bayesian weighted Mendelian randomization method) [15]. Note that those existing methods were developed for settings where a polygenic trait is analyzed as an exposure for other complex diseases and so many independent genetic variants associated with the exposure trait are available as candidate IVs. Those methods may not be suited for our target settings in which gene expression levels is considered as the exposure, and there are often only a limited number of correlated cis-eQTLs as IVs (trans-eQTLs are not considered as IVs in our two-sample MR analysis because trans-eQTL effects are less replicable across eQTL and GWAS samples). Some of those existing methods also do not allow the IVs to be correlated. Nonetheless, we included the methods for comparison. None of the existing methods were developed for taking multi-tissue eQTLs (multiple sets of IV-to-exposure association statistics) as input and that is an innovation of our method.

#### Data generation

In each simulation scenario, we simulated data for a total of *N* = *N*_*g*_ + *N*_*R*_ = 10, 300 independent subjects: *N*_*g*_ = 10, 000 subjects in a GWAS study, and *N*_*R*_ = 300 subjects in a reference multi-tissue eQTL study of *K* = 10 tissues.

First, we simulated an *N* ×*I* genotype matrix **L** for each gene, comprised of *Q* independent LD blocks with 20 SNPs in each block (thus, a total of *I* = 20 × *Q* SNPs for each gene). The correlation between SNP index *i* and SNP index *j* in a given LD block is *r*_*ij*_ = 0.95^|*i*−*j*|^, with the minor allele frequency (MAF) of SNP *i*, MAF_*i*_ ∼ Unif(0.05, 0.5). From each LD block, we randomly selected 1 SNP to be the true eQTL. The *N*_*g*_ × *Q* genotype matrix of the *Q* true eSNPs in the GWAS study is denoted **G**. For *M* (*M* ≥ 0) LD blocks, we randomly selected 1 SNP to be an invalid IV having a direct effect on the complex trait (the value of *M* varies across simulation scenarios). The *N*_*g*_ × *M* genotype matrix of the *M* SNPs that are invalid IVs is denoted **H**. We generated phenotypes in the GWAS study according to the following data generation models:

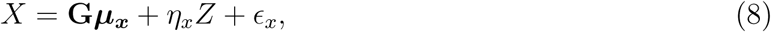

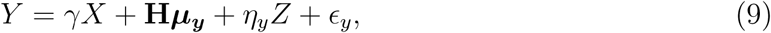

In Model (8), *X* is a vector of gene expression levels; **G** are the genotypes of eSNPs; 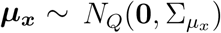 are the eQTL effects of eSNPs from independent LD blocks, with 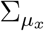 a diagonal matrix with diagonal elements 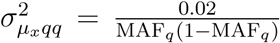; *Z* ∼ *N* (0, 1) is a vector of a latent confounder; *η*_*x*_ ∼ Unif(0, 0.1) is the effect of the confounder on gene expression levels; and *ϵ*_*x*_ ∼ *N* (0, 1) are error terms. In Model (9), *Y* is a vector of a continuous complex trait value; *γ* is the parameter of interest, the effect of gene *X* on trait *Y*, with *γ* = 0 under the null and *γ* = 0.3 under the alternative; **H** are the genotypes of SNPs having a direct effect on *Y* not through gene expression of 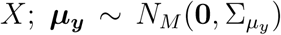 are the direct effects on *Y* of *M* SNPs from independent LD blocks, with 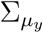 a diagonal matrix with diagonal elements 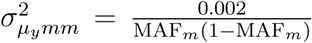; *η* ∼ Unif (0, 0.1)is the effect of the confounder on the complex trait; and *ϵ*_*y*_ ∼ *N* (0, 1) are the error terms. Across scenarios we vary *M*, the number of LD blocks having an invalid IV.

Data from the eQTL study was generated based on the model:

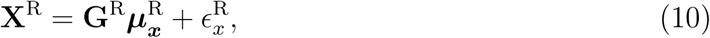

where **X**^R^ is an *N*_*R*_ × *K* matrix of expression levels measured in *K* tissues; **G**^R^ is a *N*_*R*_ × *Q* genotype matrix of *Q* eSNPs in the eQTL study; 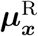 is a *Q* × *K* matrix of the tissue-specific eQTL effects; and 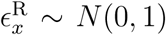 are the error terms. Each column of 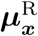 is independently drawn from *N*_*Q*_(***µ***_***x***_, 0.02 *·* **I**), where ***µ***_***x***_ is from Model (8).

After individual-level data were generated in each simulation, we calculated the marginal eQTL and GWAS summary statistics. For two-sample MR analyses, we then obtained the marginal effect estimate of each SNP *i* on gene expression in tissue *k* in the reference eQTL study, 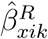; and obtained the marginal effect estimate of each SNP *i* on its simulated trait in the GWAS study, 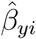. We also obtained the standard error estimates for marginal eQTL and GWAS effects.

#### Results of simulation studies

In Scenario 1, we evaluated the robustness of MR-Robin to the proportion of invalid IVs compared to existing two-sample MR methods. *P* < 0.05 was used as the significance criterion for each method. Table 1 shows the type I error rate and power comparison in the presence of 0, 10, …, 50% invalid IVs, allowing IVs to be correlated (pairwise LD *r*^2^ < 0.5 or 0.3) over 10,000 simulations of *Q* = 10 LD blocks. Since our method allows for correlated IVs and it is hard to define invalid versus valid IVs when SNPs are correlated, the proportions of invalid IVs in the tables are the proportion of LD blocks with pleiotropy, and is only an approximation of the invalid IVs among all selected ones. In each table, we also presented the average numbers of selected IVs that are from valid versus invalid LD blocks. For the competing methods, which were not developed for multi-tissue eQTL datasets, we used the eQTL summary statistics from one randomly selected tissue as input for the IV-exposure summary statistics. As shown in the table, whereas competing methods are unable to control the type I error rate when there are any invalid instruments and instruments are in LD, MR-Robin maintains reasonable control of the type I error rate if a majority of instruments are valid (e.g. up to 30% invalid IVs). The last three methods in the table were developed for independent instruments; since they do not account for correlation (LD) among the instruments, they do not control the type I error rate even when all instruments are valid. Power is reasonable for all methods when a majority of IVs are valid. In Supplemental Materials, Tables S1-2, we compared the type I error rates and powers using alternative LD selection criteria for the IVs (pairwise LD *r*^2^ < 0.1 or 0.01).

**Table 1:** Simulation results evaluating the performance of MR-Robin. Averaged type I error rates and power over 10,000 simulations are shown by percentage of invalid instruments (using *P* < 0.05 as the significance criterion for each method). 10 LD blocks were simulated, with one true eQTL per LD block. Instruments were selected sequentially: the eSNP with the strongest association with gene expression was selected, and the next selected eSNP is the strongest-associated SNP remaining also with LD *r*^2^ < *ρ* with any already-selected eSNPs. Results shown for *ρ* = 0.5 (A) and *ρ* = 0.3 (B)

| Method | Proportion of invalid IV (%) |  |  |  |  |  |
| --- | --- | --- | --- | --- | --- | --- |
|  | 0 | 10 | 20 | 30 | 40 | 50 |
|  | Type I error rate |  |  |  |  |  |
| MR-Robin | 0.047 | 0.051 | 0.059 | 0.061 | 0.071 | 0.085 |
| MR-Egger | 0.032 | 0.057 | 0.087 | 0.110 | 0.122 | 0.138 |
| MR-RAPS | 0.255 | 0.303 | 0.348 | 0.380 | 0.399 | 0.426 |
| MRMix | 0.136 | 0.175 | 0.207 | 0.223 | 0.256 | 0.267 |
| BWMR | 0.279 | 0.349 | 0.396 | 0.440 | 0.452 | 0.479 |
|  | Power |  |  |  |  |  |
| MR-Robin | 0.800 | 0.763 | 0.725 | 0.685 | 0.659 | 0.615 |
| MR-Egger | 0.942 | 0.931 | 0.923 | 0.917 | 0.914 | 0.905 |
| MR-RAPS | 0.996 | 0.993 | 0.986 | 0.979 | 0.976 | 0.962 |
| MRMix | 0.511 | 0.499 | 0.498 | 0.502 | 0.491 | 0.493 |
| BWMR | 0.998 | 0.994 | 0.988 | 0.981 | 0.978 | 0.966 |
|  | Avg number of SNPs selected (valid/invalid) |  |  |  |  |  |
| All Methods | 30.4 /0.0 | 27.4 /3.0 | 24.4 /6.0 | 21.4 /9.2 | 18.3 /12.1 | 15.2 /15.2 |

(B) pairwise LD $r^2 < 0.3$
| Method | Proportion of invalid IV (%) |  |  |  |  |  |
| --- | --- | --- | --- | --- | --- | --- |
|  | 0 | 10 | 20 | 30 | 40 | 50 |
|  | Type I error rate |  |  |  |  |  |
| MR-Robin | 0.047 | 0.051 | 0.055 | 0.051 | 0.061 | 0.062 |
| MR-Egger | 0.027 | 0.062 | 0.097 | 0.126 | 0.137 | 0.161 |
| MR-RAPS | 0.084 | 0.118 | 0.147 | 0.170 | 0.186 | 0.214 |
| MRMix | 0.130 | 0.199 | 0.237 | 0.273 | 0.288 | 0.307 |
| BWMR | 0.097 | 0.141 | 0.182 | 0.206 | 0.222 | 0.246 |
|  | Power |  |  |  |  |  |
| MR-Robin | 0.679 | 0.640 | 0.605 | 0.577 | 0.549 | 0.505 |
| MR-Egger | 0.894 | 0.886 | 0.878 | 0.870 | 0.860 | 0.853 |
| MR-RAPS | 0.986 | 0.979 | 0.969 | 0.955 | 0.944 | 0.925 |
| MRMix | 0.512 | 0.515 | 0.517 | 0.509 | 0.500 | 0.498 |
| BWMR | 0.992 | 0.985 | 0.975 | 0.962 | 0.952 | 0.936 |
|  | Avg number of SNPs selected (valid/invalid) |  |  |  |  |  |
| All Methods | 14.1 /0.0 | 12.7 /1.4 | 11.3 /2.8 | 9.9 /4.2 | 8.5 /5.6 | 7.0 /7.0 |

In the second simulation scenario, we evaluated the performance of MR-Robin when the number of selected IVs is small. We simulated the data using *Q* = 3 LD blocks, with two blocks without pleiotropy and one block with pleiotropy (thus the proportion of LD blocks with pleiotropic effects is fixed at 33.3%). Table 2 shows the type I error rates and power when the selection LD *r*^2^ threshold is set to 0.5, 0.3, 0.2, 0.1 and 0.01. As shown in the table, MR-Robin performs reasonably well even when the number of IVs is very limited. Though in this setting, MR-Robin requires the IVs to be less dependent (*r*^2^ < 0.3). MR-Robin outperforms competing methods in this setting.

**Table 2:** Simulation results evaluating the performance of MR-Robin when there is a small number of IVs. Averaged type I error rates and power over 10,000 simulations are shown by IV selection criteria. 3 LD blocks were simulated, with two blocks without pleiotropic effects (valid IVs) and one block with (invalid IV). Results shown for five IV selection criteria (LD *r*^2^ < 0.5, 0.3, 0.2, 0.1, and 0.01).

| Method | LD selection criteria ($r^2$) | | | | |
| --- | --- | --- | --- | --- | --- |
|  | 0.5 | 0.3 | 0.2 | 0.1 | 0.01 |
|  | Type I error rate |  |  |  |  |
| MR-Robin | 0.084 | 0.056 | 0.048 | 0.039 | 0.032 |
| MR-Egger | 0.087 | 0.100 | 0.111 | 0.139 | 0.158 |
| MR-RAPS | 0.380 | 0.203 | 0.144 | 0.104 | 0.091 |
| MRMix | 0.254 | 0.240 | 0.239 | 0.227 | 0.223 |
| BWMR | 0.431 | 0.227 | 0.154 | 0.103 | 0.087 |
|  | Power |  |  |  |  |
| MR-Robin | 0.586 | 0.455 | 0.391 | 0.320 | 0.292 |
| MR-Egger | 0.676 | 0.625 | 0.602 | 0.553 | 0.504 |
| MR-RAPS | 0.860 | 0.799 | 0.773 | 0.740 | 0.726 |
| MRMix | 0.473 | 0.470 | 0.480 | 0.487 | 0.496 |
| BWMR | 0.884 | 0.811 | 0.769 | 0.722 | 0.695 |
|  | Avg # of SNPs selected |  |  |  |  |
| All Methods | 7.4 /3.7 | 3.5 /1.8 | 2.7 /1.4 | 2.2 /1.1 | 2.0 /1.0 |

The simulation results showed that MR-Robin is able to control the type I error using correlated instruments provided that a majority (≥ 70%) of the instruments are valid IVs. Moreover, in Table 2, we showed that even when the number of available IVs is very small (3-10), the proposed MR-Robin can still yield reasonable results if the small number of IVs are relatively less dependent (*r*^2^ < 0.3). Last but not least, we want to emphasize that when IVs are correlated, if one IV is an invalid IV, all the other correlated IVs are also affected to some degree, and as such the random-slope model of MR-Robin with its resampling-based inference procedure fits the need for allowing correlated IVs when considering the effect of gene expression on a complex trait.

### Application: Identifying schizophrenia (SCZ) risk-associated genes via MR-Robin

To detect genes with expression levels being associated with schizophrenia risk, we applied MR-Robin using summary statistics from two-samples: schizophrenia risk GWAS statistics from the second schizophrenia mega-analysis (SCZ2) conducted by the Psychiatric Genomics Consortium (PGC) [34], and multi-tissue eQTL statistics from the 13 brain tissues in version 8 (V8) of the Genotype-Tissue Expression (GTEx) project [33]. Details of the two datasets can be found in Supplemental Materials.

We first formed the set of instrumental variables (IVs) for each gene by selecting the cis-eSNPs/IVs (within 1 Mb of transcription start site) and the brain tissue types in which they have strong IV effects. All the cis-SNPs being selected are cross-tissue IVs (with median eQTL *P* < 0.05). However, it is well known in the IV literature that weak IVs, i.e., SNPs being only weakly associated with the genes, would result in high variance and misleading inferences even when they are valid IVs [35; 36]. And therefore, we will choose cross-tissue eQTLs with significant eQTL effects of *P* -value ≤ 0.001 in at least three tissue types, i.e., being reasonably strong IVs to provide reliable inferences [37] in at least three tissue types. And we restrict the analysis to the cross-tissue IVs in the tissue types with strong cross-tissue (or shared) effects. Since this step involves only the selection of IV based on the strength of the eQTL effects, with no information regarding the outcome, the selection of IV and tissue types would not induce inflation in false positive findings.

While the analysis is restricted to strong IVs with *P* < 0.001 in at least 3 tissues, we iteratively selected the (next) best eSNP satisfying the IV selection criteria and having pairwise LD *r*^2^ < 0.5 with each of the eSNPs already selected. Note that here we conducted the primary analysis with a relatively liberal LD threshold to improve the power of the analysis. Following the primary analysis, we later conducted a sensitivity analysis on the implied genes to check the robustness of our results to the choice of IVs. If the gene has only 1 cis-eQTL, MR-Robin would be reduced to a single-IV analysis, which can be heavily affected by the validity of the IV with assumptions that cannot be adequately checked in general. Therefore, we restricted the MR-Robin analysis to 3,127 protein-coding genes with at least 5 IVs selected based on this criteria. For each SNP/IV used in the analysis, we used eQTL statistics only from those brain tissues where the SNP had eQTL *P* < 0.001 (with strong IV effects). Thus, each SNP/IV has 3-13 observed eQTL effect estimates from different tissue types in the unbalanced mixed effects model.

At a false discovery rate (FDR) < 5%, we identified 43 genes as showing evidence of a dependence between gene expression levels and SCZ risk. For the 43 genes whose expression showed an association with SCZ risk in the primary analysis, we performed a sensitivity analysis using different IV selection criteria. Specifically in the sensitivity analysis, for each gene, among the cross-tissue IVs with median eQTL *P* < 0.05 having strong IV effects in at least 3 tissues (*P* < 0.001), we iteratively dropped the eSNP with the highest correlation to others until all pairwise LD *r*^2^ < 0.3 among remaining eSNPs or only 5 eSNPs remained. For each eSNP, we still only used eQTL statistics from tissues where that eSNP had eQTL *P* < 0.001. In the sensitivity analysis, there were 39 and 42 genes with MR-Robin *P* < 0.05 and *P* < 0.1, respectively, all of which had a fixed effect estimate matching the sign of the fixed effect estimate from the primary analysis.

Figure 2 plotted the multi-tissue eQTL effect sizes in the GTEx brain tissues against the GWAS effect sizes in the PGC dataset for two selected genes in the primary analysis (left column) versus the sensitivity analysis (right column). The gene *THOC7* (Figure 2A) showed consistent correlations between eQTL and GWAS effects based on two sets of correlated Ivs in the primary and sensitivity analyses (both with *P* < 5 × 10^−3^). Despite some SNPs having a potentially larger deviation from the shared effect than the others – indicated by the random slopes (colored lines) deviating from the fixed effect estimate (black line) – the plot shows a clear pattern of association between the magnitude of eQTL effects and magnitude of GWAS effects, implying that the expression levels of *THOC7* affect schizophrenia risk. The protein encoded by *THOC7* is a component of the THO complex of the TRanscription and EXport (TREX) complex which couples transcription to mRNA export, specifically associating with spliced mRNA [38; 39]. Mutations in subunits of TREX have been associated with neurodevelopmental disorders [40], and a recent TWAS study that imputed gene expression in brain tissues found an association between expression levels of *THOC7* in cerebellum and schizophrenia risk [41]. In contrast, the gene *RNF149* (Figure 2B) was the only gene no longer significant in the sensitivity analysis (*P* = 0.15), and prompts further exploration. The change in significance for *RNF149* may be at least partially due to an increase in the relative proportion among selected IVs that have potential pleiotropic effects (i.e. better fitted by a line with non-zero intercept in Figure 2B) when using more stringent LD *r*^2^ selection criteria. In the Supplemental Materials, we presented additional details and the scatterplots of multi-tissue eQTL effect estimates against SCZ GWAS effect estimates for selected IVs of all 42 genes identified by MR-Robin in the primary analysis having *P* < 0.1 the sensitivity analyses.

**Figure 2:**
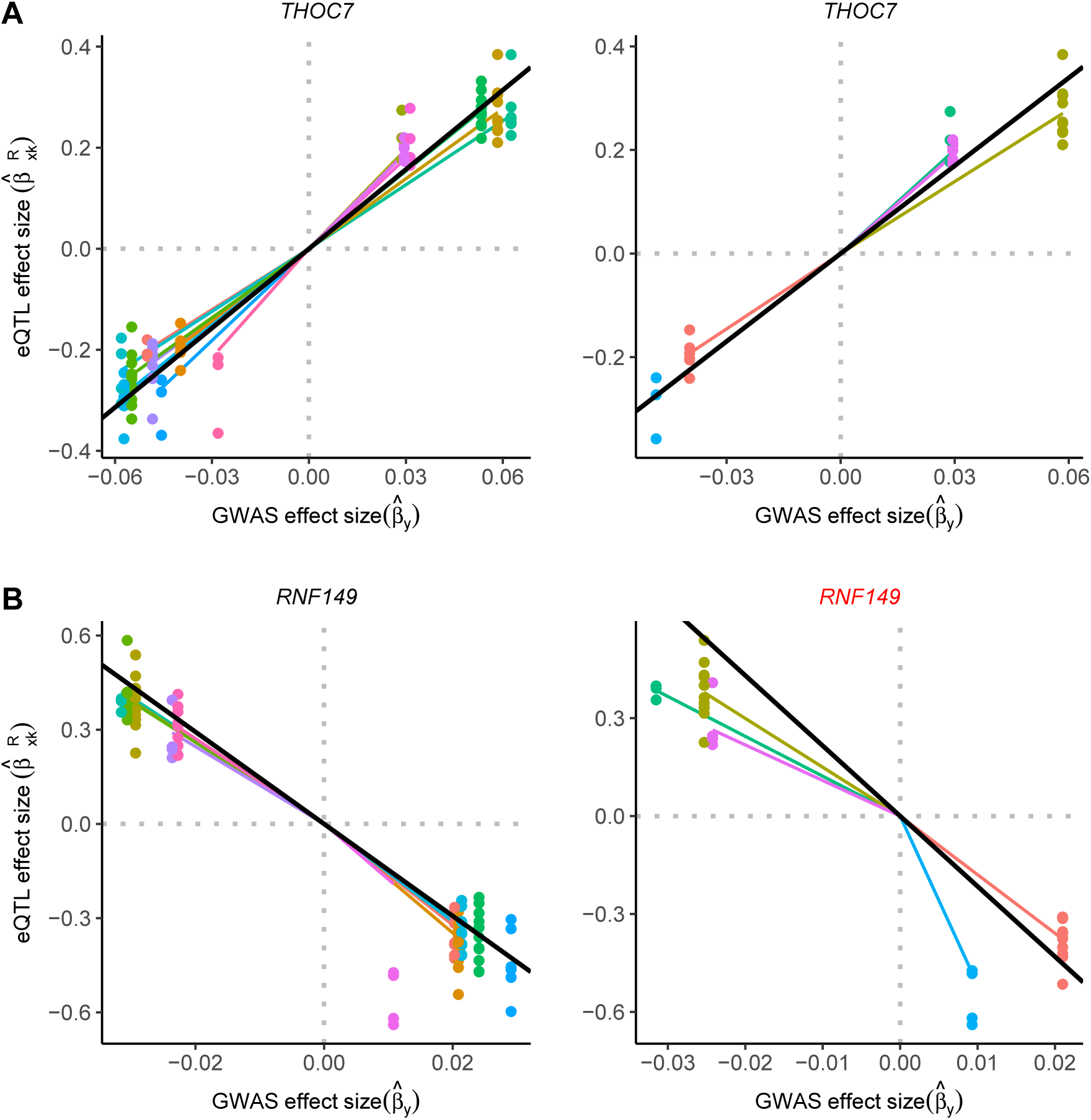
Illustrations of two example genes in the primary analysis (left column) and the sensitivity analysis (right column). Multi-tissue eQTL effect sizes in the GTEx brain tissues were plotted against SCZ GWAS effect sizes in the PGC dataset for the genes, *THOC7* and *RNF149*. In the sensitivity analysis using alternative IV selection criteria, the SCZ risk association remained for *THOC7* (A) (*P* < 0.1) with consistency in the sign of the fixed effect estimate. The association was no longer significant between *RNF149* expression (B) and SCZ risk in the sensitivity analysis (*P* = 0.15). Points are colored by SNP. Colored lines represent SNP-specific slope estimates. The slope of the black line is the fixed effect estimate from the MR-Robin reverse regression. The results imply a non-zero effect of the gene *THOC7* on schizophrenia risk.

In summary, we applied the newly proposed two-sample MR method, MR-Robin, to integrate multiple sets of brain tissue eQTL summary statistics from GTEx and SCZ GWAS summary statistics from PGC, and have identified 42 genes with potential causal associations to schizophrenia risk. These 42 genes demonstrated consistent dependencies between brain eQTL and SCZ GWAS association effects using two sets of SNPs as IVs based on different selection criteria. The results highlighted the value of MR-Robin as a robust two-sample MR method that allows moderately correlated and some invalid instrumental variables and identifies gene expression levels as causal exposures for complex diseases.

## Discussion

In this work, we proposed a robust two-sample MR method – MR-Robin – allowing correlated and invalid IVs. MR-Robin was motivated by analyses of gene expression levels as causal exposures for complex diseases/traits. In those settings, often only a limited number of potentially correlated cis-eQTLs are available as candidate instrumental variables (IVs), posing new challenges to MR analyses. MR-Robin integrates GWAS statistics with multitissue eQTL statistics in a mixed model framework, considering the estimated effect of gene expression levels on disease from each IV as an observed value of the true effect plus a SNP-specific bias. Compared to existing robust two-sample MR methods, a major innovation of MR-Robin is the use of multi-tissue eQTL summary statistics (multiple sets of IV-to-exposure statistics). Based on a reverse regression framework with multi-tissue eQTL effects as response, the rich information in multi-tissue eQTL data allows the estimation of SNP-specific random slopes (due to being in LD with SNPs with horizontal and/or correlated pleiotropy) as well as the fixed-effects correlation of eQTL and GWAS effects across all IVs based on a limited number of IVs. In contrast, existing models and methods based on the deconvolution of mixture distributions or penalized regressions in general require a large number of IVs to achieve stability in estimation. To account for correlation among IVs due to LD and tissue-tissue correlations, MR-Robin utilizes a resampling procedure when testing the effect from gene expression levels to the complex trait. We showed through simulations that MR-Robin was able to control the type I error rates using a limited number of moderately correlated IVs when the proportion of IVs that are invalid is moderate.

We applied MR-Robin to identify genes with expression levels affecting schizophrenia risk by integrating multiple sets of brain tissue eQTL statistics from GTEx and SCZ GWAS statistics from PGC. We identified 42 genes showing consistent dependencies between multi-tissue eQTL and GWAS association effects based on two different sets of IVs with different selection criteria from primary and sensitivity analyses. Our analysis illustrated that MR-Robin and two-sample MR methods, requiring only multi-tissue eQTL and GWAS summary statistics as input, could be used as another integrative method in recapitalizing on existing summary statistics to further map gene expression levels or other omics traits affecting a complex trait of interest, to explain the potential mechanisms underlying trait susceptibility loci, and to identify clinically actionable targets with larger effects on complex diseases and traits.

There are several caveats and limitations to the current work. First, similar to other two-sample MR methods, MR-Robin cannot by itself prove a causal relationship from a gene to a complex trait but rather suggests instances consistent with a causal model. Nevertheless, analyses using MR-Robin may be useful in prioritizing candidate genes for additional follow-up and research. Second, MR-Robin requires summary statistics from a multi-tissue eQTL dataset as input. For some complex traits being considered as the outcome, it may not be obvious which tissues are most relevant to the trait being studied. Several recent works have proposed methods or provided resources to identify trait-relevant tissues [42; 43; 44], and these works may be useful in such cases. Third, to accurately estimate the SNP-specific bias, MR-Robin requires more than one SNP to be used as an IV. Depending on the dataset and IV selection criteria, there may be some genes whose association with the complex trait cannot be appropriately tested using MR-Robin.

MR-Robin was developed as a two-sample MR method to test for effects from the expression levels of a gene on a complex trait. MR-Robin can be applied to discover genes that may be causally associated with a complex trait of interest or to confirm that a putative gene demonstrates consistency with a model in which its gene expression causally affects the complex trait. The method may also be extended more generally to settings where a limited number of potentially correlated candidate IVs are present provided that multiple estimates of either the IV-exposure or IV-outcome statistics are available.

The R package MrRobin is available at https://github.com/kjgleason/MrRobin.

## Acknowledgements

We thank the GTEx Consortium. We thank Drs. Francois Aguet, Kristin Ardlie for providing cis-eQTL summary statistics. We thank Drs. Jiebiao Wang, Jubao Duan, Xin He and Brandon L. Pierce for providing valuable suggestions to the development of the methods and analysis.

## Funding

This work was supported by the National Institutes of Health (NIH) grant R01GM108711 and 2R01GM108711 to LSC and KJG. LSC was also supported by SUB-U24 CA2109993. KJG was also supported by Susan G. Komen ® GTDR16376189 and the National Cancer Institute of the NIH under Award Number F31CA239557. FY was supported by the NIH grant R03CA208387 and 2R01GM108711.

## Availability of data and material

The R package MrRobin is available at https://github.com/kjgleason/MrRobin.

## Competing interests

The authors declare that they have no competing interests.

## Notes

### Competing Interest Statement

The authors have declared no competing interest.

