## Supplemental Materials for "A robust two-sample Mendelian Randomization method integrating GWAS with multi-tissue eQTL summary statistics"

June 5, 2020

#### Additional simulations to evaluate the performance of MR-Robin compared to existing methods

In the main text, we have presented the performance of MR-Robin when the selected IVs were in moderate LD ($r^2 < 0.5$ and $r^2 < 0.3$). In this section, using simulation studies we evaluated the performance of MR-Robin when the selected IVs are in weak LD or are nearly independent, and compared to competing methods. In each simulation scenario, we simulated data for a total of $N = N_g + N_R = 10,300$ independent subjects: $N_g = 10,000$ subjects in a GWAS study, and $N_R = 300$ subjects in a reference multitissue eQTL study of $K = 10$ tissues. Details of the simulations are described in the main text.

#### MR-Robin controls type I error rate with moderate proportion of invalid IVs

In Scenario 1, we evaluated the robustness of MR-Robin to the proportion of invalid IVs. We simulated the data using $Q = 10$ LD blocks with 20 SNPs in each, varying the proportion of invalid IVs across settings. That is, we varied the proportion of LD blocks in which a SNP has direct effects on the complex trait $Y$ (i.e. effects not mediated through gene expression $X$). Over 10,000 simulations, we compare the type I error rate and power of MR-Robin to existing two-sample MR methods. $P < 0.05$ was used as the significance criterion for each method. Tables S1A and S1B compare the methods when the selection LD $r^2$ threshold is set to 0.1 and 0.01, respectively (results using selection LD $r^2$ thresholds of 0.5 and 0.3 are reported in Table 1 in main text).

Based on the results, we observe that competing methods are generally unable to control the type I error rate when there are a moderate proportion of invalid IVs (e.g $> 20\%$) and IVs are in weak LD. This is mostly because their estimating algorithm may require a relatively large number of IVs to perform well and in our target settings there are often only limited number of independent cis-eQTLs in a region. On the other hand, MR-Robin is able to control the type I error rate when a majority of IVs are valid (with up to 50% invalid IVs) if the selected IVs are in weak LD. See Table 1 in the main text for simulation results with alternative LD selection criteria (i.e. LD $r^2$ thresholds of 0.5 and 0.3) and Table 2 in the main text for simulation results when the number of candidate IVs is very small (i.e. $Q = 3$ LD blocks each with 20 SNPs).

Since our method allows for correlated IVs and it is hard to define invalid versus valid IVs when SNPs are correlated, the proportions of invalid IVs in the tables are the proportion of LD blocks with pleiotropy, and is only an approximation of the invalid IVs among all selected ones. In each table, we also presented the average numbers of selected IVs that are from valid versus invalid LD blocks.

Table S1: Simulation results evaluating the performance of MR-Robin. Averaged type I error rates and power over 10,000 simulations are shown by percentage of invalid instruments. 10 LD blocks were simulated, with one true eQTL per LD block. Instruments were selected sequentially: the eSNP with the strongest association with gene expression was selected, and the next selected eSNP is the strongest-associated SNP remaining also with LD $r^2 < 0.1$ (A) or $r^2 < 0.01$ (B) with any already-selected eSNPs.

(A) pairwise LD $r^2 < 0.1$

| Method | Proportion of invalid IV (%) | | | | | |
|---|---|---|---|---|---|---|
| | 0 | 10 | 20 | 30 | 40 | 50 |
| | Type I error rate | | | | | |
| MR-Robin | 0.052 | 0.049 | 0.047 | 0.047 | 0.054 | 0.049 |
| MR-Egger | 0.023 | 0.078 | 0.129 | 0.162 | 0.190 | 0.221 |
| MR-RAPS | 0.030 | 0.051 | 0.061 | 0.074 | 0.090 | 0.100 |
| MRMix | 0.113 | 0.190 | 0.240 | 0.275 | 0.301 | 0.313 |
| BWMR | 0.037 | 0.059 | 0.073 | 0.086 | 0.101 | 0.106 |
| | Power | | | | | |
| MR-Robin | 0.588 | 0.553 | 0.522 | 0.493 | 0.484 | 0.435 |
| MR-Egger | 0.809 | 0.798 | 0.786 | 0.778 | 0.767 | 0.762 |
| MR-RAPS | 0.962 | 0.950 | 0.937 | 0.919 | 0.903 | 0.878 |
| MRMix | 0.564 | 0.563 | 0.550 | 0.545 | 0.544 | 0.542 |
| BWMR | 0.975 | 0.961 | 0.947 | 0.931 | 0.914 | 0.889 |
| | Avg number of SNPs selected (valid/invalid) | | | | | |
| All Methods | 7.8 /0.0 | 7.1 /0.8 | 6.3 /1.6 | 5.5 /2.4 | 4.7 /3.1 | 3.9 /3.9 |

(B) pairwise LD $r^2 < 0.01$

| Method | Proportion of invalid IV (%) | | | | | |
|---|---|---|---|---|---|---|
| | 0 | 10 | 20 | 30 | 40 | 50 |
| | Type I error rate | | | | | |
| MR-Robin | 0.052 | 0.049 | 0.050 | 0.047 | 0.048 | 0.045 |
| MR-Egger | 0.020 | 0.077 | 0.127 | 0.170 | 0.192 | 0.236 |
| MR-RAPS | 0.024 | 0.045 | 0.058 | 0.072 | 0.086 | 0.096 |
| MRMix | 0.106 | 0.182 | 0.230 | 0.263 | 0.305 | 0.318 |
| BWMR | 0.031 | 0.052 | 0.066 | 0.078 | 0.088 | 0.098 |
| | Power | | | | | |
| MR-Robin | 0.650 | 0.608 | 0.573 | 0.541 | 0.523 | 0.484 |
| MR-Egger | 0.770 | 0.754 | 0.740 | 0.733 | 0.725 | 0.717 |
| MR-RAPS | 0.957 | 0.947 | 0.931 | 0.915 | 0.898 | 0.876 |
| MRMix | 0.590 | 0.586 | 0.580 | 0.572 | 0.578 | 0.566 |
| BWMR | 0.969 | 0.954 | 0.936 | 0.925 | 0.902 | 0.880 |
| | Avg number of SNPs selected (valid/invalid) | | | | | |
| All Methods | 6.3 /0.0 | 5.7 /0.6 | 5.0 /1.3 | 4.4 /1.9 | 3.8 /2.5 | 3.1 /3.1 |

### MR-Robin identified genes showing evidence of association with schizophrenia risk

In Table S2, we present detailed information on the 42 genes identified by MR-Robin as being consistent with a causal model in which gene expression affects schizophrenia risk. These 42 genes were identified using a false discovery rate (FDR) [S1] threshold of $< 5\%$ in the primary analysis and maintained their association (MR-Robin $P < 0.1$) in the sensitivity analysis. For a given IV-gene pair, only summary statistics from tissues with eQTL $P < 0.001$ were used in the MR-Robin analysis.

Scatterplots plotting multi-tissue eQTL effect sizes in the GTEx brain tissues against the GWAS effect sizes in the PGC dataset for selected IVs of the 42 identified genes are shown in Figure S1 (following the Supplemental References). Despite some SNPs having a potentially larger deviation from the shared effect than the others – indicated by the random slopes (colored lines) deviating from the fixed effect estimate (black line) – the plots generally show clear patterns of association between the magnitude of eQTL effects and magnitude of GWAS effects, implying that the expression levels of these genes affect schizophrenia risk.

Table S2: Detailed information on the 42 schizophrenia risk-associated genes identified by MR-Robin at FDR < 5% in the primary analysis and also having MR-Robin $P < 0.1$ in the sensitivity analysis. IVs were selected from cross-tissue eSNPs (median eQTL $P < 0.05$) that were strong IVs ($P < 0.001$) in at least 3 tissues. In the primary analysis, the strongest eSNP having pairwise LD $r^2 < 0.5$ was iteratively selected. In the sensitivity analysis, the eSNP with the highest pairwise correlation with other SNPs was iteratively removed until pairwise LD $r^2 < 0.3$ among all eSNPs or only 5 eSNPs remained. For a given IV, only summary statistics from tissues with eQTL $P < 0.001$ were used in the MR-Robin analysis.

| Gene info | | | MR-Robin $P$-values | |
|---|---|---|---|---|
| Ensembl ID | Gene Symbol | Chromosome | Primary | Sensitivity |
| ENSG00000048544 | *MRPS10* | 6 | $4.2 \times 10^{-4}$ | $3.5 \times 10^{-2}$ |
| ENSG00000089486 | *CDIP1* | 16 | $4.5 \times 10^{-6}$ | $4.2 \times 10^{-4}$ |
| ENSG00000090263 | *MRPS33* | 7 | $2.7 \times 10^{-4}$ | $1.6 \times 10^{-3}$ |
| ENSG00000100731 | *PCNX1* | 14 | $1.6 \times 10^{-5}$ | $1.5 \times 10^{-4}$ |
| ENSG00000115649 | *CNPPD1* | 2 | $4.6 \times 10^{-5}$ | $4.3 \times 10^{-5}$ |
| ENSG00000117601 | *SERPINC1* | 1 | $5.7 \times 10^{-4}$ | $1.2 \times 10^{-2}$ |
| ENSG00000120451 | *SNX19* | 11 | $4.7 \times 10^{-5}$ | $5.0 \times 10^{-4}$ |
| ENSG00000123643 | *SLC36A1* | 5 | $9.7 \times 10^{-4}$ | $7.7 \times 10^{-3}$ |
| ENSG00000125611 | *CHCHD5* | 2 | $4.8 \times 10^{-4}$ | $6.0 \times 10^{-2}$ |
| ENSG00000126464 | *PRR12* | 19 | $2.9 \times 10^{-4}$ | $7.0 \times 10^{-3}$ |
| ENSG00000129925 | *TMEM8A* | 16 | $< 1.0 \times 10^{-6}$ | $< 1.0 \times 10^{-6}$ |
| ENSG00000130304 | *SLC27A1* | 19 | $1.5 \times 10^{-4}$ | $1.4 \times 10^{-3}$ |
| ENSG00000141013 | *GAS8* | 16 | $4.0 \times 10^{-4}$ | $3.7 \times 10^{-3}$ |
| ENSG00000141127 | *PRPSAP2* | 17 | $7.0 \times 10^{-4}$ | $6.5 \times 10^{-2}$ |
| ENSG00000142233 | *NTN5* | 19 | $4.5 \times 10^{-5}$ | $4.3 \times 10^{-5}$ |
| ENSG00000142534 | *RPS11* | 19 | $3.0 \times 10^{-4}$ | $2.1 \times 10^{-4}$ |
| ENSG00000142599 | *RERE* | 1 | $2.8 \times 10^{-4}$ | $6.7 \times 10^{-4}$ |
| ENSG00000146966 | *DENND2A* | 7 | $< 1.0 \times 10^{-6}$ | $< 1.0 \times 10^{-6}$ |
| ENSG00000147403 | *RPL10* | X | $2.7 \times 10^{-4}$ | $3.3 \times 10^{-3}$ |
| ENSG00000159199 | *ATP5G1* | 17 | $6.4 \times 10^{-4}$ | $3.5 \times 10^{-3}$ |
| ENSG00000162753 | *SLC9C2* | 1 | $5.2 \times 10^{-6}$ | $1.9 \times 10^{-3}$ |
| ENSG00000163040 | *CCDC74A* | 2 | $1.1 \times 10^{-4}$ | $7.1 \times 10^{-3}$ |
| ENSG00000163634 | *THOC7* | 3 | $4.7 \times 10^{-4}$ | $5.0 \times 10^{-3}$ |
| ENSG00000163938 | *GNL3* | 3 | $3.1 \times 10^{-6}$ | $7.2 \times 10^{-4}$ |
| ENSG00000167468 | *GPX4* | 19 | $3.2 \times 10^{-4}$ | $9.6 \times 10^{-2}$ |
| ENSG00000167535 | *CACNB3* | 12 | $< 1.0 \times 10^{-6}$ | $4.1 \times 10^{-5}$ |
| ENSG00000169220 | *RGS14* | 5 | $6.3 \times 10^{-5}$ | $5.3 \times 10^{-4}$ |
| ENSG00000170802 | *FOXN2* | 2 | $1.0 \times 10^{-4}$ | $2.9 \times 10^{-4}$ |
| ENSG00000171928 | *TVP23B* | 17 | $7.2 \times 10^{-4}$ | $1.7 \times 10^{-4}$ |
| ENSG00000173273 | *TNKS* | 8 | $6.7 \times 10^{-4}$ | $4.2 \times 10^{-3}$ |
| ENSG00000177595 | *PIDD1* | 11 | $2.1 \times 10^{-4}$ | $1.9 \times 10^{-3}$ |
| ENSG00000177707 | *NECTIN3* | 3 | $1.6 \times 10^{-4}$ | $7.0 \times 10^{-4}$ |
| ENSG00000180921 | *FAM83H* | 8 | $4.4 \times 10^{-4}$ | $9.2 \times 10^{-3}$ |
| ENSG00000182093 | *WRB* | 21 | $< 1.0 \times 10^{-6}$ | $< 1.0 \times 10^{-6}$ |
| ENSG00000196268 | *ZNF493* | 19 | $< 1.0 \times 10^{-6}$ | $7.1 \times 10^{-6}$ |
| ENSG00000203499 | *FAM83H-AS1* | 8 | $8.8 \times 10^{-4}$ | $3.6 \times 10^{-3}$ |
| ENSG00000204257 | *HLA-DMA* | 6 | $< 1.0 \times 10^{-6}$ | $3.1 \times 10^{-6}$ |
| ENSG00000204962 | *PCDHA8* | 5 | $9.9 \times 10^{-4}$ | $8.0 \times 10^{-3}$ |
| ENSG00000214013 | *GANC* | 15 | $3.3 \times 10^{-5}$ | $4.5 \times 10^{-3}$ |
| ENSG00000224389 | *C4B* | 6 | $7.8 \times 10^{-4}$ | $2.3 \times 10^{-2}$ |
| ENSG00000225190 | *PLEKHM1* | 17 | $5.2 \times 10^{-5}$ | $4.0 \times 10^{-5}$ |
| ENSG00000244731 | *C4A* | 6 | $4.5 \times 10^{-5}$ | $1.7 \times 10^{-4}$ |

### Description of data used in analyses

#### The Genotype-Tissue Expression project (GTEx)

The Genotype-Tissue Expression (GTEx) project is building a comprehensive resource to study tissue-specific gene expression and regulation by collecting post-mortem tissue samples from non-diseased tissue sites [S2]. Data analyzed in this paper is from GTEx version 8 (V8) [S3]. GTEx samples underwent Whole Genome Sequencing to a median depth of 32X on Illumina HiSeq 2000 or Illumina HiSeq X. GTEx RNA sequencing was performed using the Illumina TruSeq™ RNA Sequencing platform. Raw sequence data were processed using the Broad's Picard pipeline [S4]. Data was aligned using STAR (v2.5.3a) [S5]. RNA-SeQC [S6] was used for quality control and gene-level expression quantification, and TMM [S7] was used to normalize read counts. Additional details about the genotyping pipeline and sample and variant quality control, and on the RNA-Sequencing pipeline and processing are reported elsewhere [S3]. Covariates adjusted for in analyses of GTEx brain tissues included gender, 5 genotype Principal Components, genotyping platform and up to 30 PEER [S8] variables.

#### The Psychiatric Genomics Consortium

Schizophrenia-risk GWAS statistics were obtained from the second schizophrenia mega-analysis (SCZ2) conducted by the Psychiatric Genomics Consortium [S9]. The GWAS was conducted using up to 36,989 cases and 113,075 controls. In the final analysis, 128 LD-independent SNPs in 108 loci were reported as surpassing the genome-wide significance threshold ($P < 5 \times 10^{-8}$). Additional details of the second PGC GWAS of schizophrenia-risk are reported elsewhere [S9].

### Reference

Figure S1: Schizophrenia risk associated genes identified by MR-Robin. Multi-tissue eQTL effect size estimates in the GTEx brain tissues (y-axis) are plotted against GWAS effect size estimates in the PGC dataset (x-axis) for SNPs used in the primary analysis (left column) and sensitivity analysis (right column). Points are colored by SNP. Colored lines represent SNP-specific slope estimates. The slope of the black line is the fixed effect estimate from the MR-Robin reverse regression.

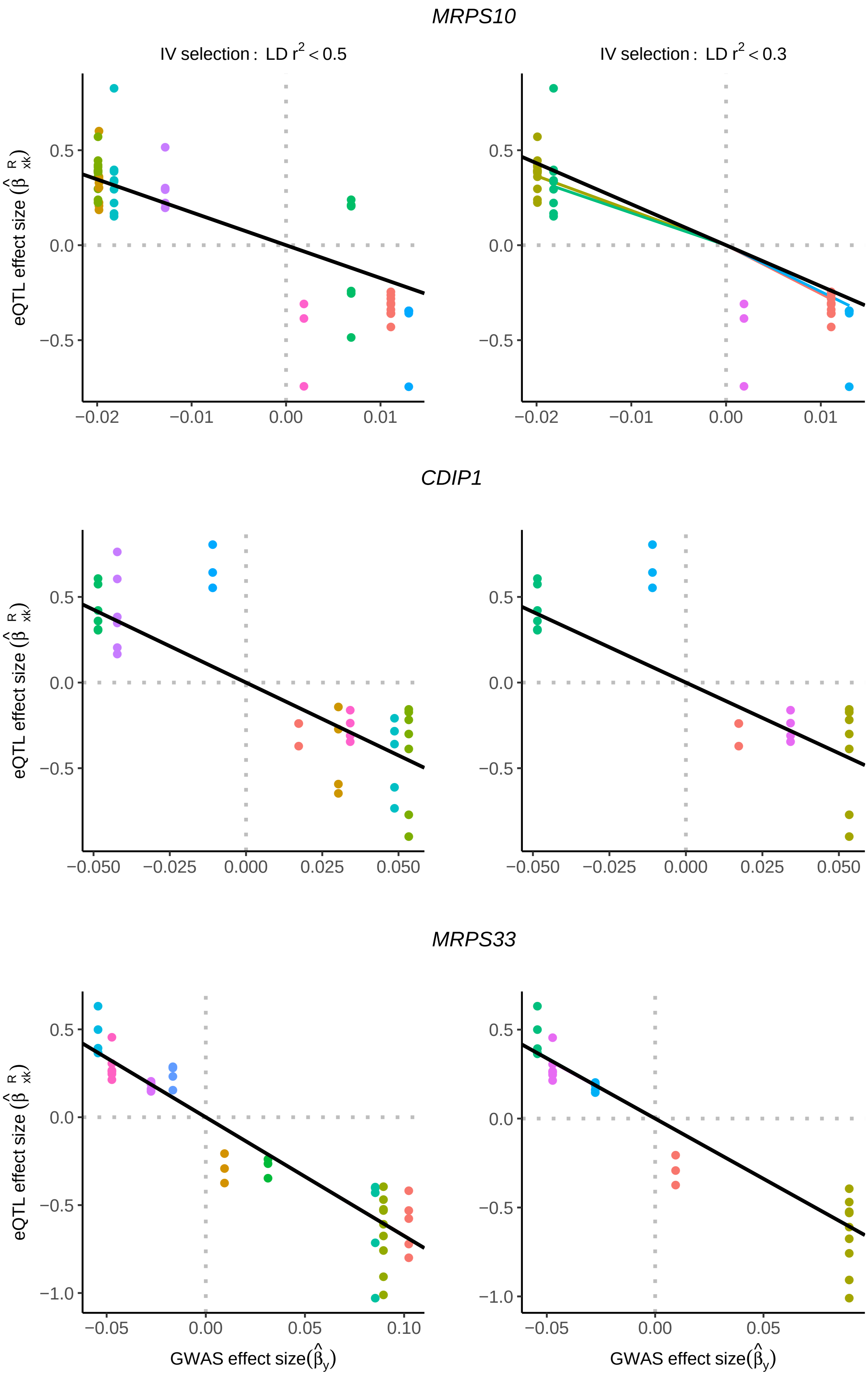
MRPS10
IV selection : LD r2 < 0.5
IV selection : LD r2 < 0.3
CDIP1
MRPS33
eQTL effect size (β̂ R xk)
GWAS effect size(β̂y)
0.5
0.0
−0.5
−1.0
−0.02
−0.01
0.00
0.01
−0.050
−0.025
0.000
0.025
0.050
−0.05
0.05
0.10

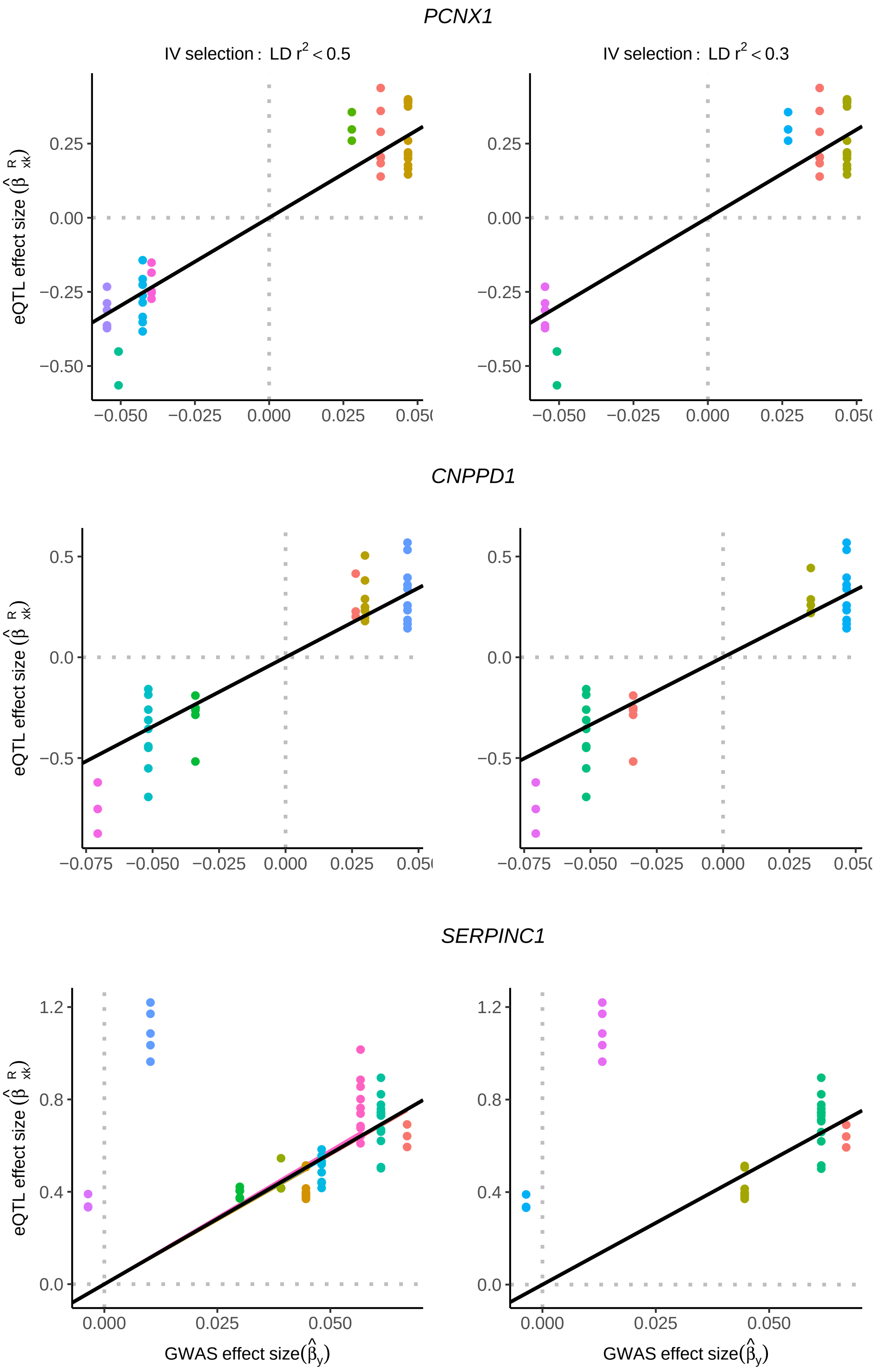
PCNX1
IV selection : LD r² < 0.5
IV selection : LD r² < 0.3
CNPPD1
SERPINC1
eQTL effect size (β̂ R xk)
GWAS effect size(β̂y)
0.25
0.00
−0.25
−0.50
0.5
0.0
−0.5
1.2
0.8
0.4
−0.075
−0.050
−0.025
0.000
0.025
0.050

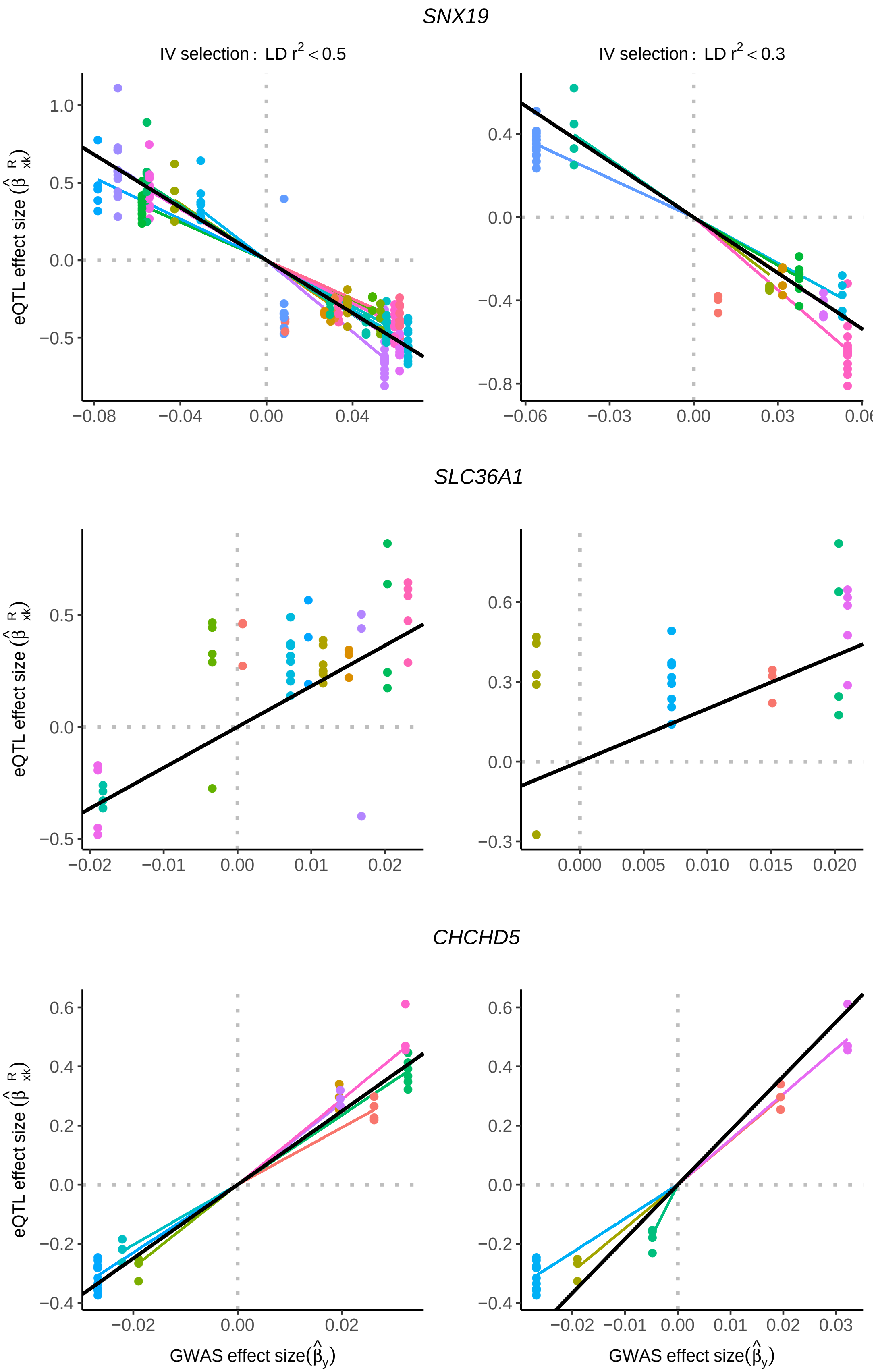
SNX19
IV selection : LD r² < 0.5
IV selection : LD r² < 0.3
eQTL effect size (β̂ R xk)
SLC36A1
CHCHD5
GWAS effect size(β̂y)

1

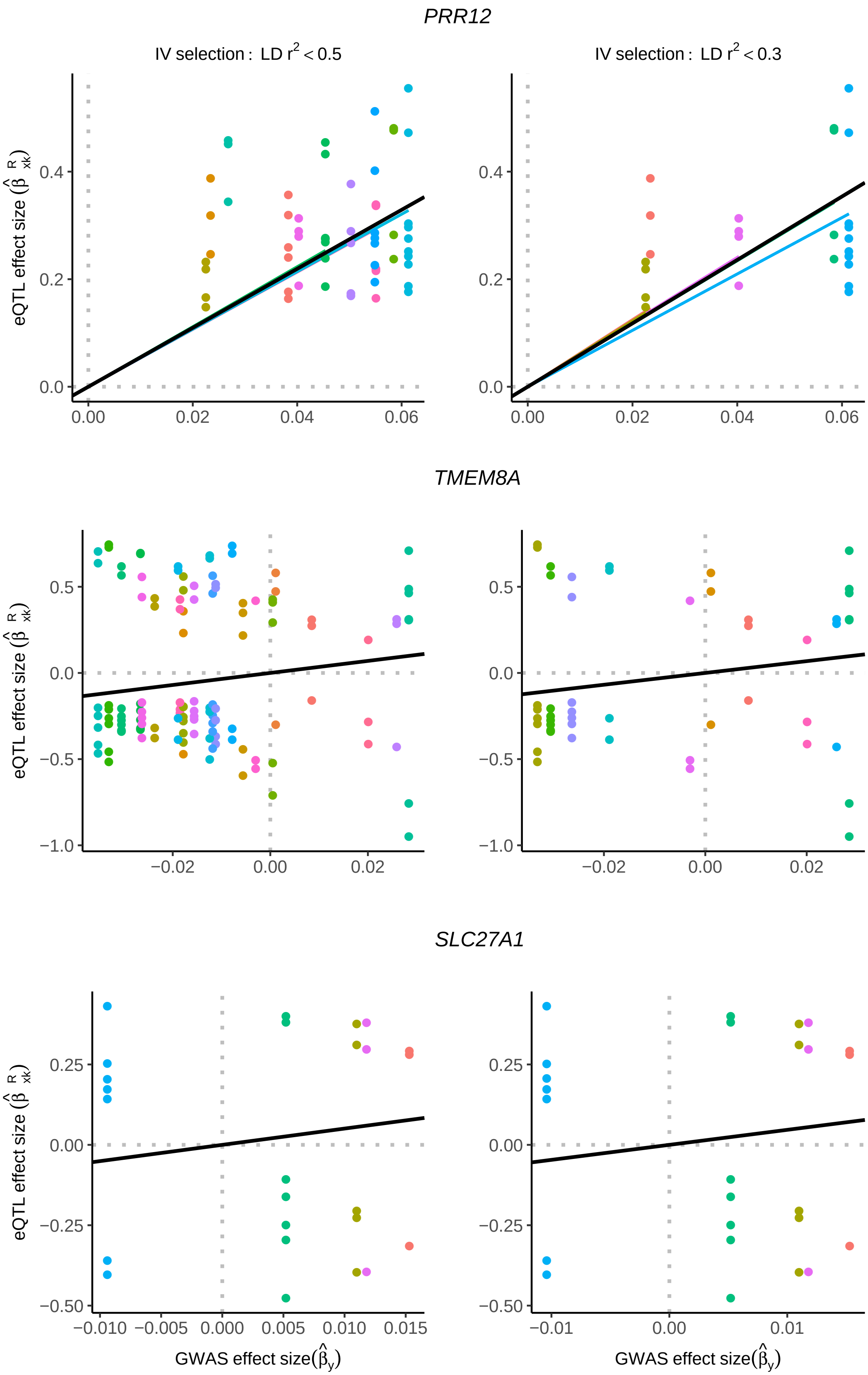

PRR12
IV selection : LD r2 < 0.5
IV selection : LD r2 < 0.3
eQTL effect size
0.0
0.2
0.4
0.00
0.02
0.04
0.06
TMEM8A
−1.0
−0.5
0.0
0.5
−0.02
0.00
0.02
SLC27A1
−0.50
−0.25
0.00
0.25
−0.010
−0.005
0.000
0.005
0.010
0.015
−0.01
0.00
0.01
GWAS effect size

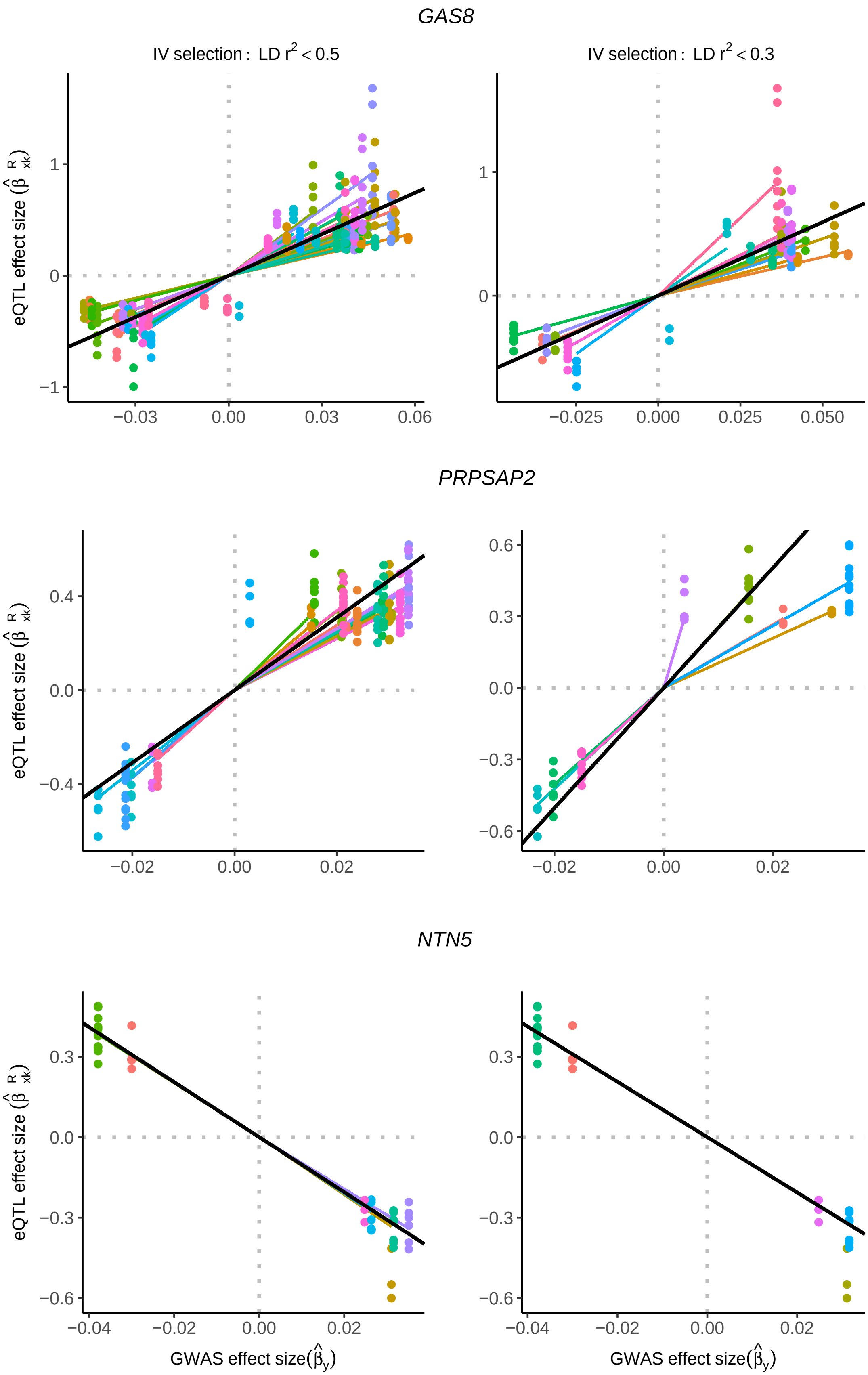
GAS8
IV selection : LD r2 < 0.5
IV selection : LD r2 < 0.3
PRPSAP2
NTN5
eQTL effect size (β̂ R xk)
GWAS effect size(β̂y)
1
0
−1
−0.03
0.00
0.03
0.06
−0.025
0.000
0.025
0.050
0.4
0.0
−0.4
0.6
0.3
−0.3
−0.6
−0.02
0.02
−0.04

1

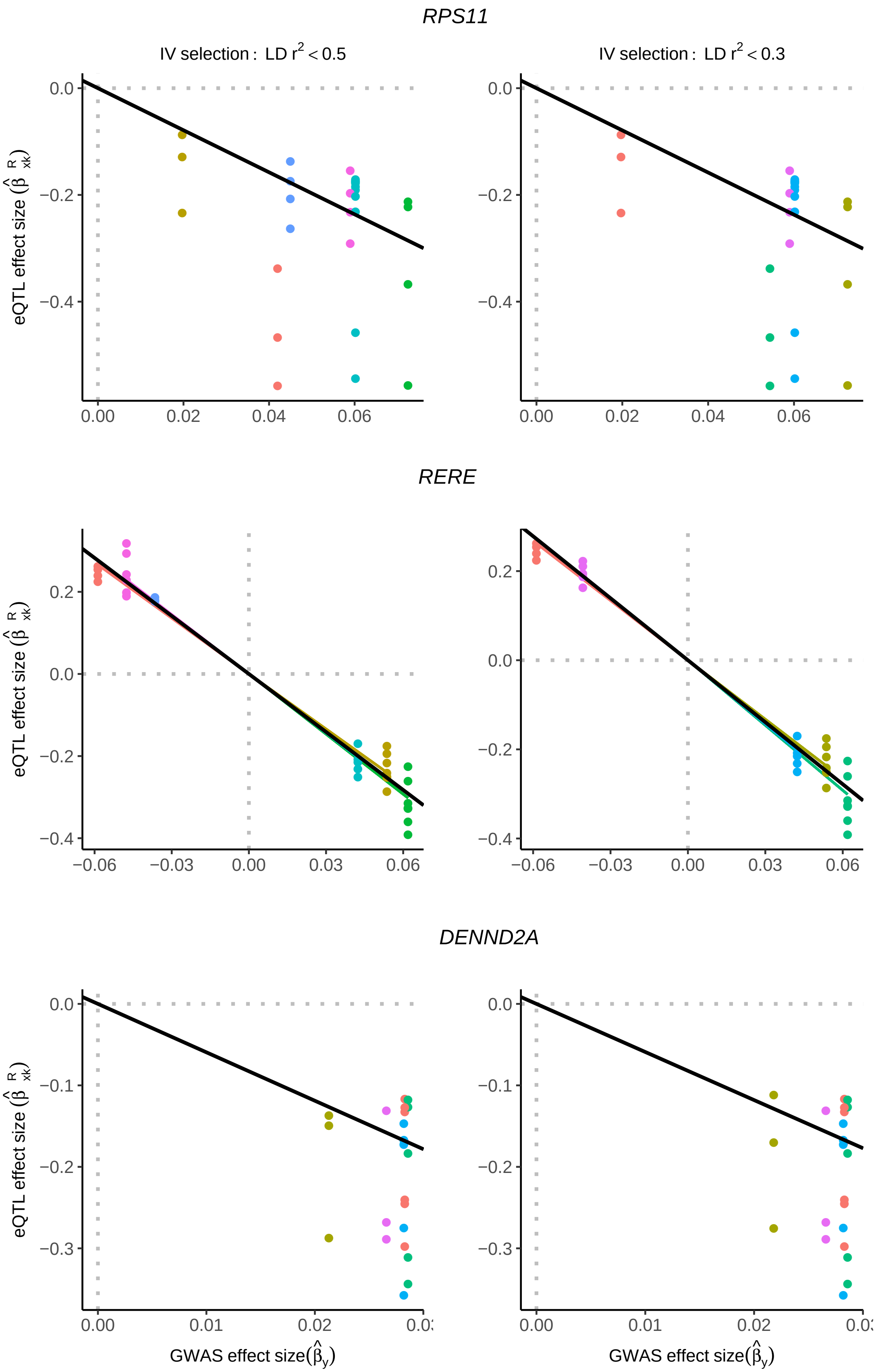

RPS11
IV selection : LD r^2 < 0.5
IV selection : LD r^2 < 0.3
eQTL effect size
RERE
DENND2A
GWAS effect size

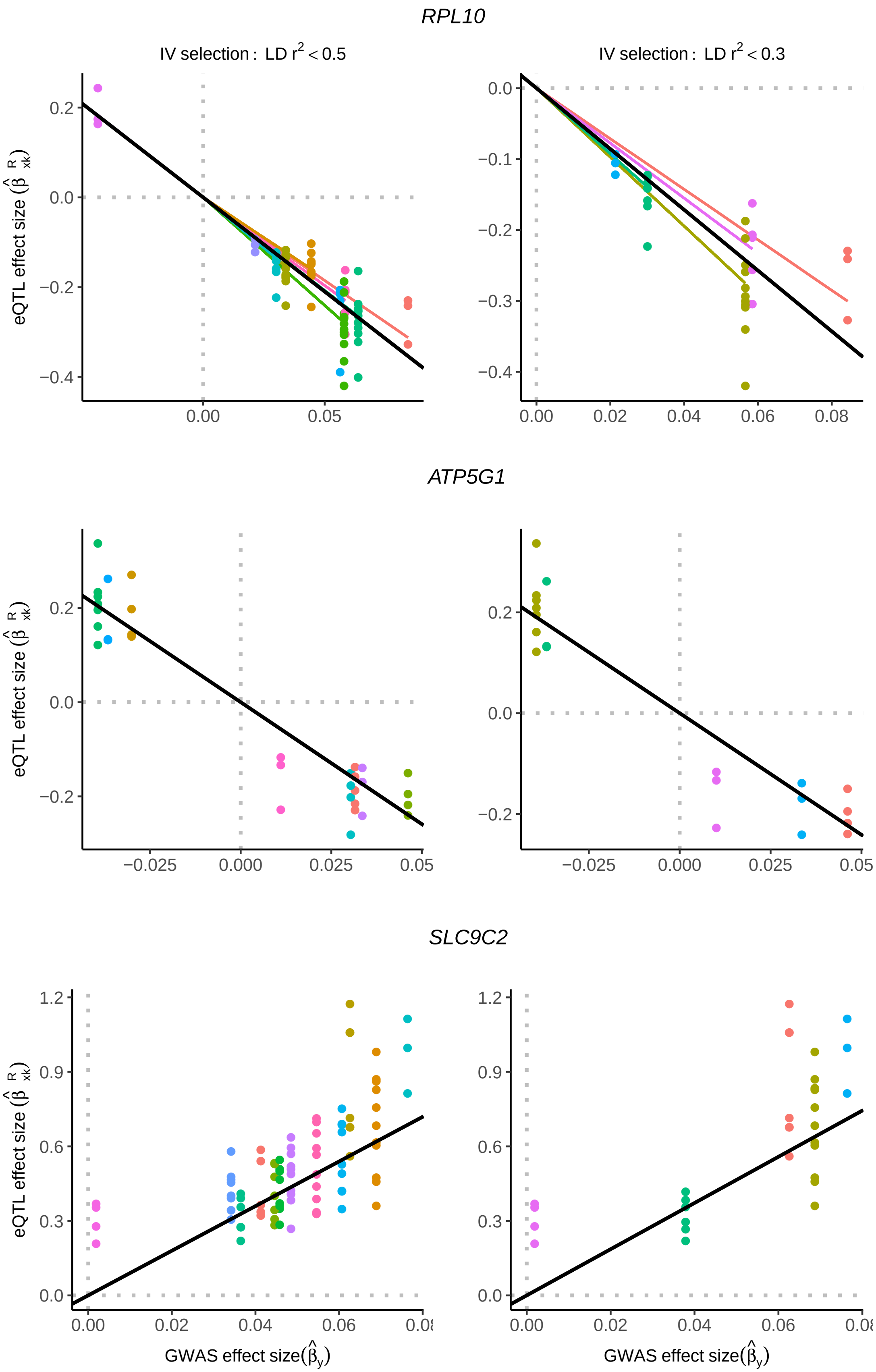

RPL10
IV selection : LD r² < 0.5
IV selection : LD r² < 0.3
ATP5G1
SLC9C2
eQTL effect size (β̂ R xk)
GWAS effect size(β̂y)

1

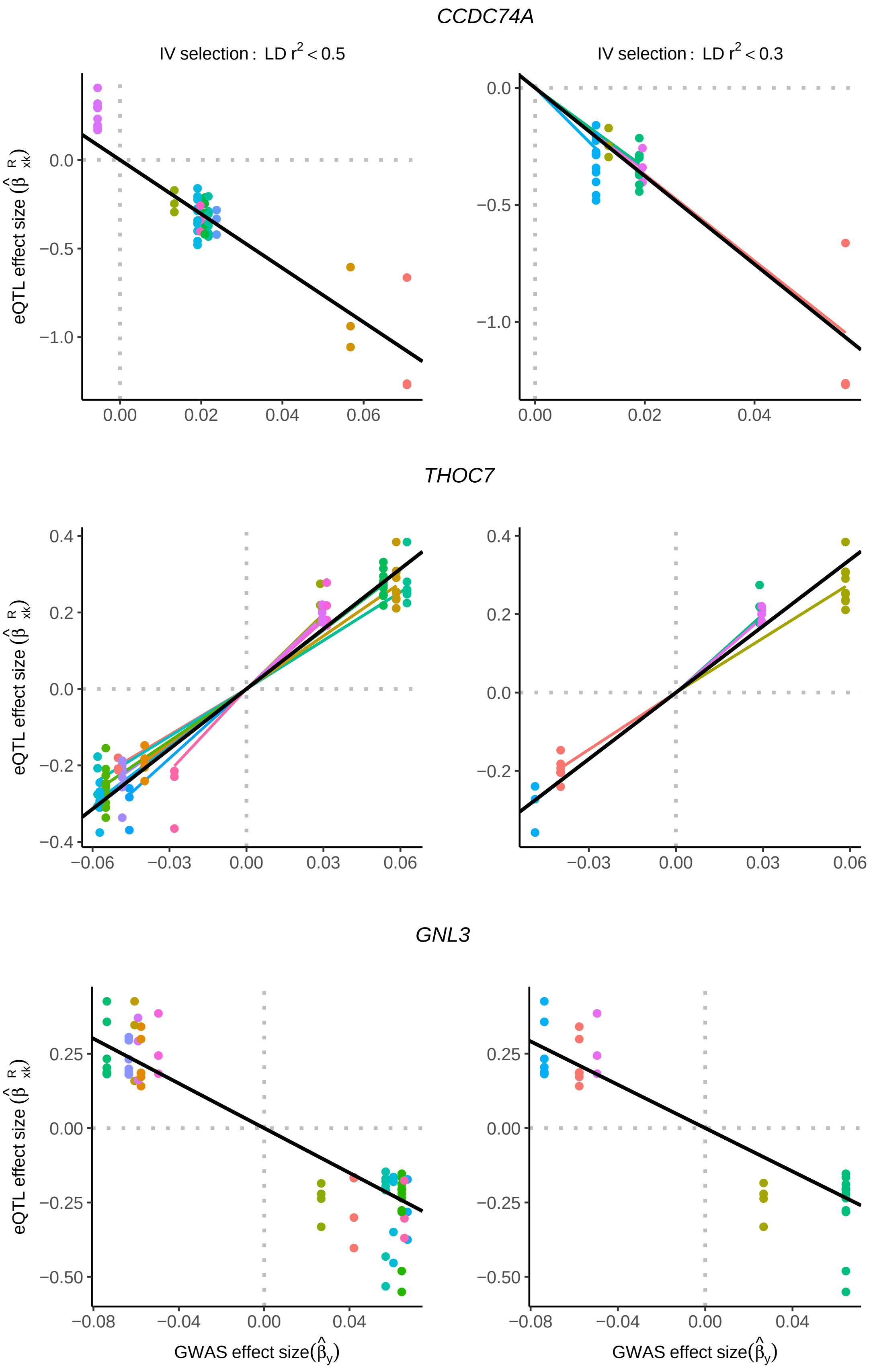
CCDC74A
IV selection : LD r² < 0.5
IV selection : LD r² < 0.3
THOC7
GNL3
eQTL effect size (β̂ R xk)
GWAS effect size(β̂y)
0.0
−0.5
−1.0
0.00
0.02
0.04
0.06
0.4
0.2
−0.2
−0.4
−0.06
−0.03
0.03
0.25
0.00
−0.25
−0.50
−0.08
−0.04

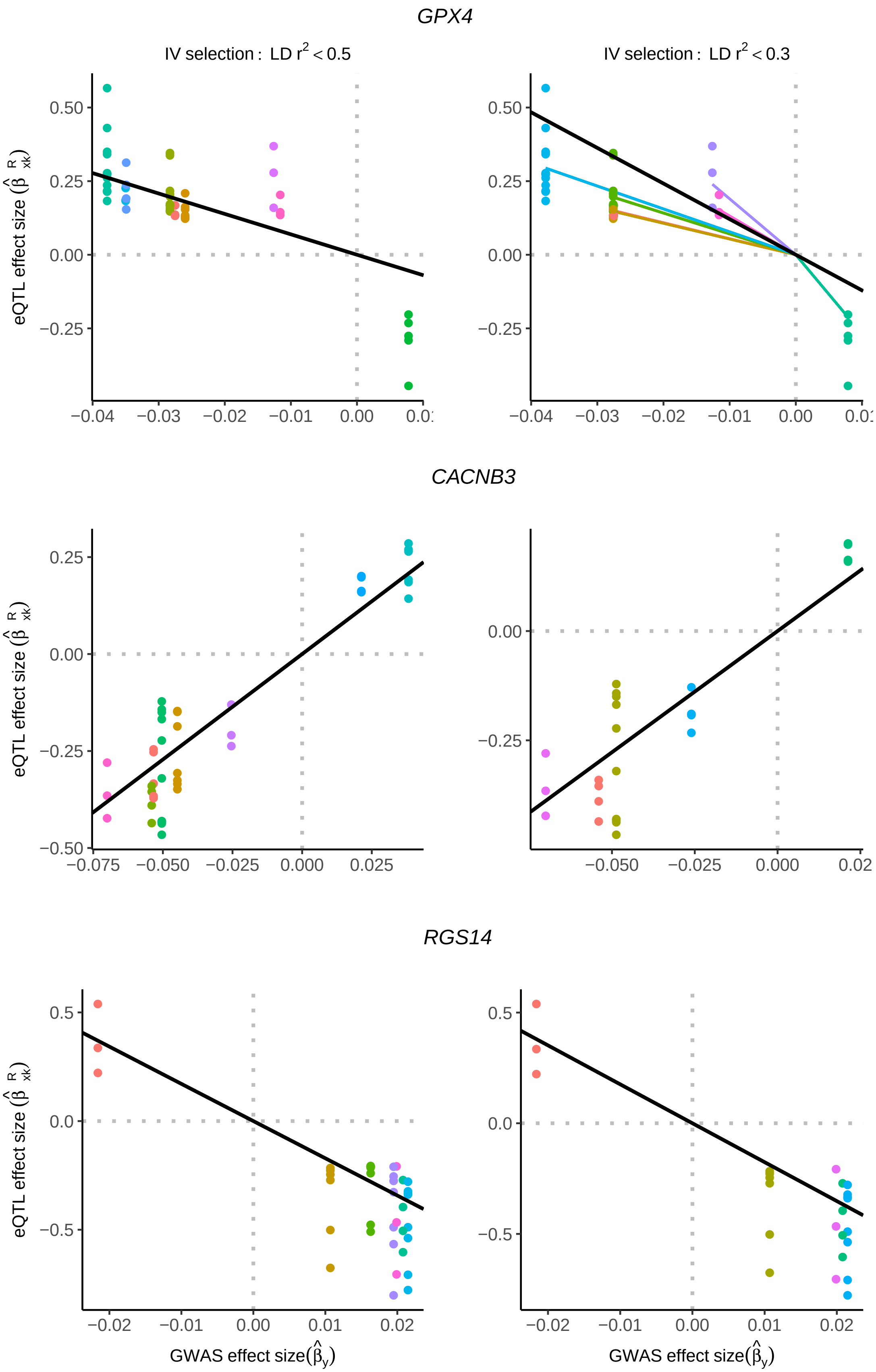

GPX4
IV selection : LD r² < 0.5
IV selection : LD r² < 0.3
CACNB3
RGS14
eQTL effect size
GWAS effect size

1

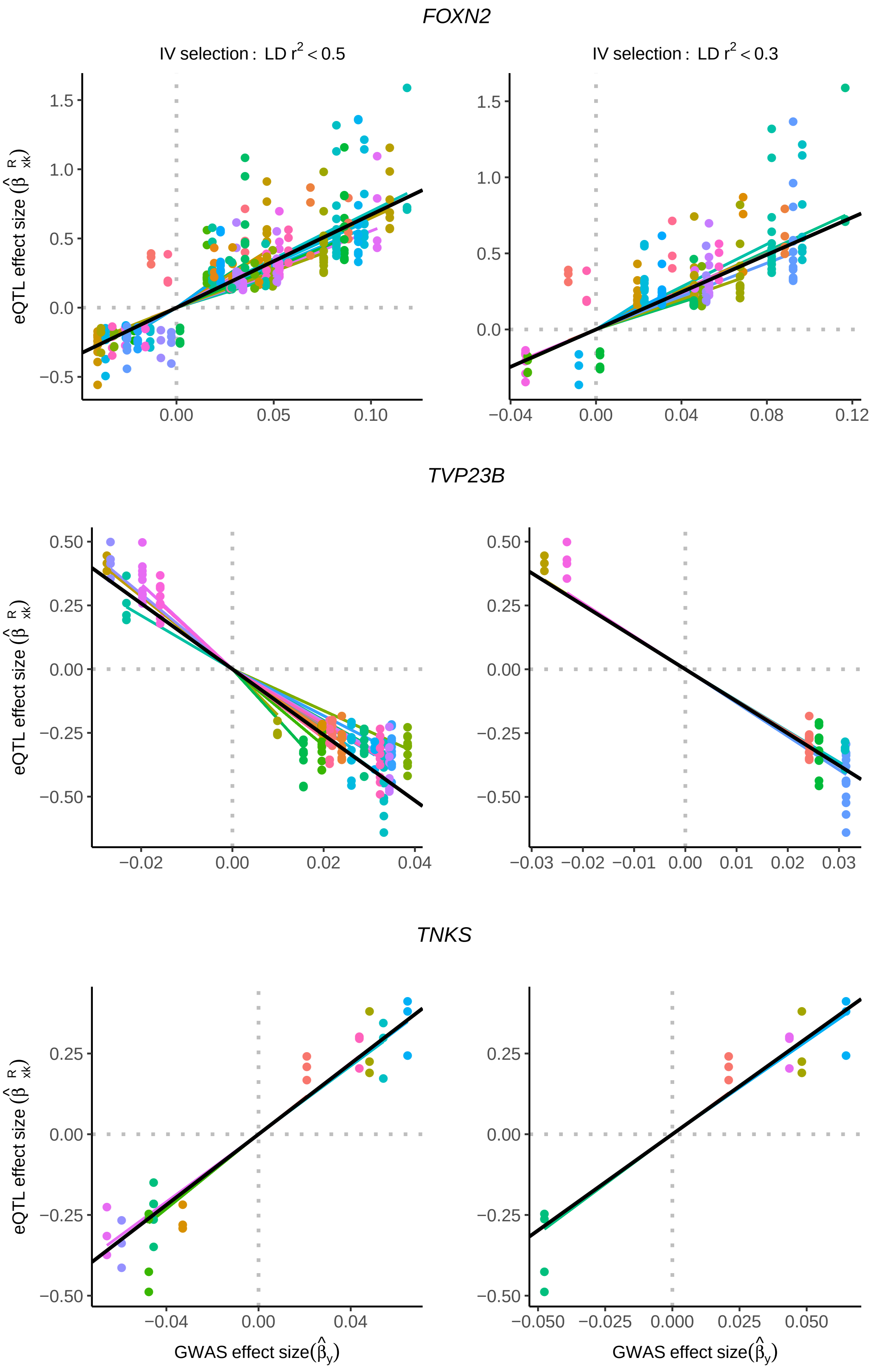

FOXN2
IV selection : LD r^2 < 0.5
IV selection : LD r^2 < 0.3
TVP23B
TNKS
eQTL effect size ($\hat{\beta}^R_{xk}$)
GWAS effect size($\hat{\beta}_y$)

1

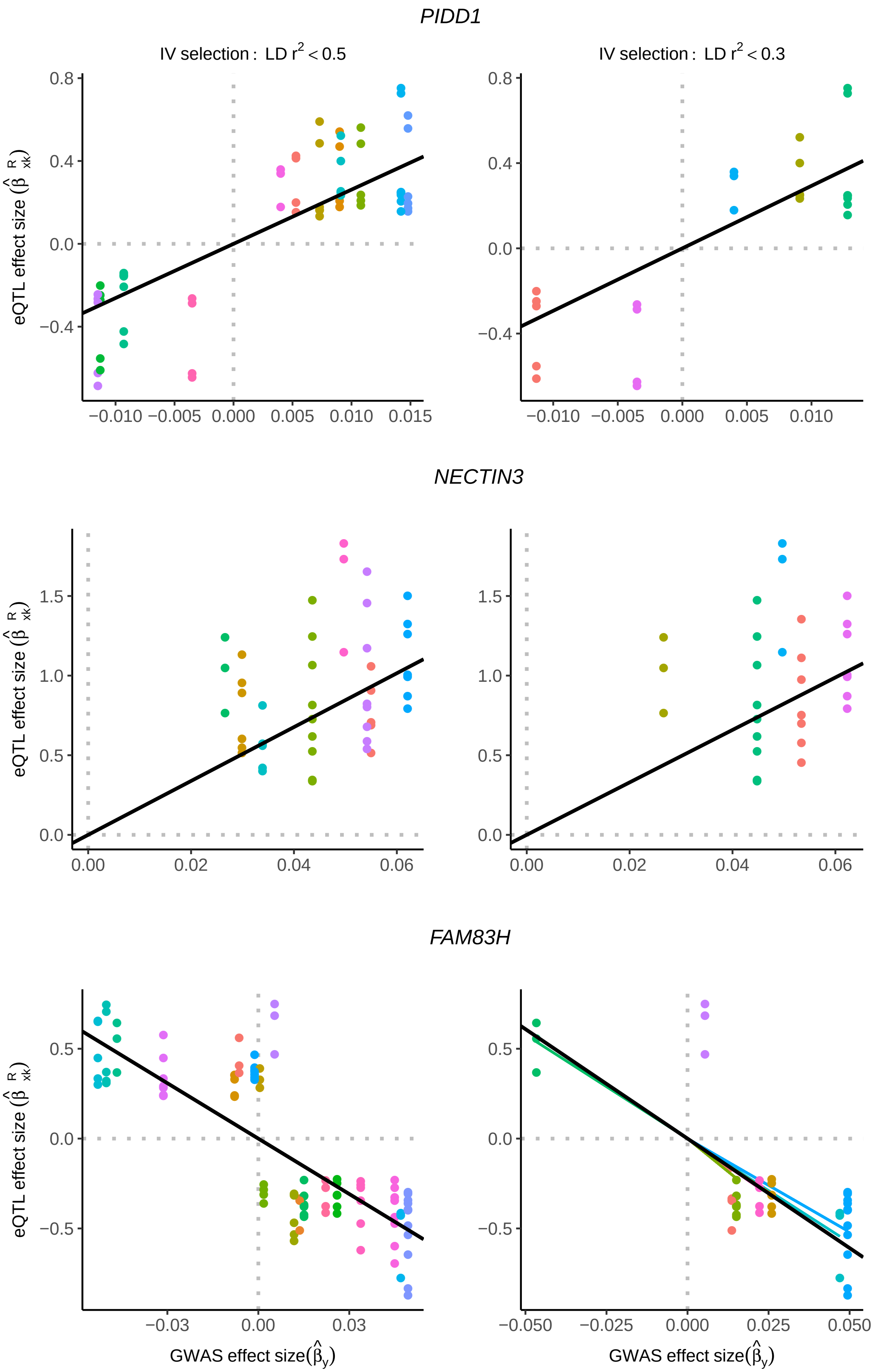
PIDD1
IV selection : LD r^2 < 0.5
IV selection : LD r^2 < 0.3
eQTL effect size (β̂ R xk)
0.8
0.4
0.0
−0.4
−0.010 −0.005 0.000 0.005 0.010 0.015
−0.010 −0.005 0.000 0.005 0.010
NECTIN3
1.5
1.0
0.5
0.0
0.00 0.02 0.04 0.06
FAM83H
−0.5
−0.03 0.00 0.03
−0.050 −0.025 0.000 0.025 0.050
GWAS effect size(β̂y)

1

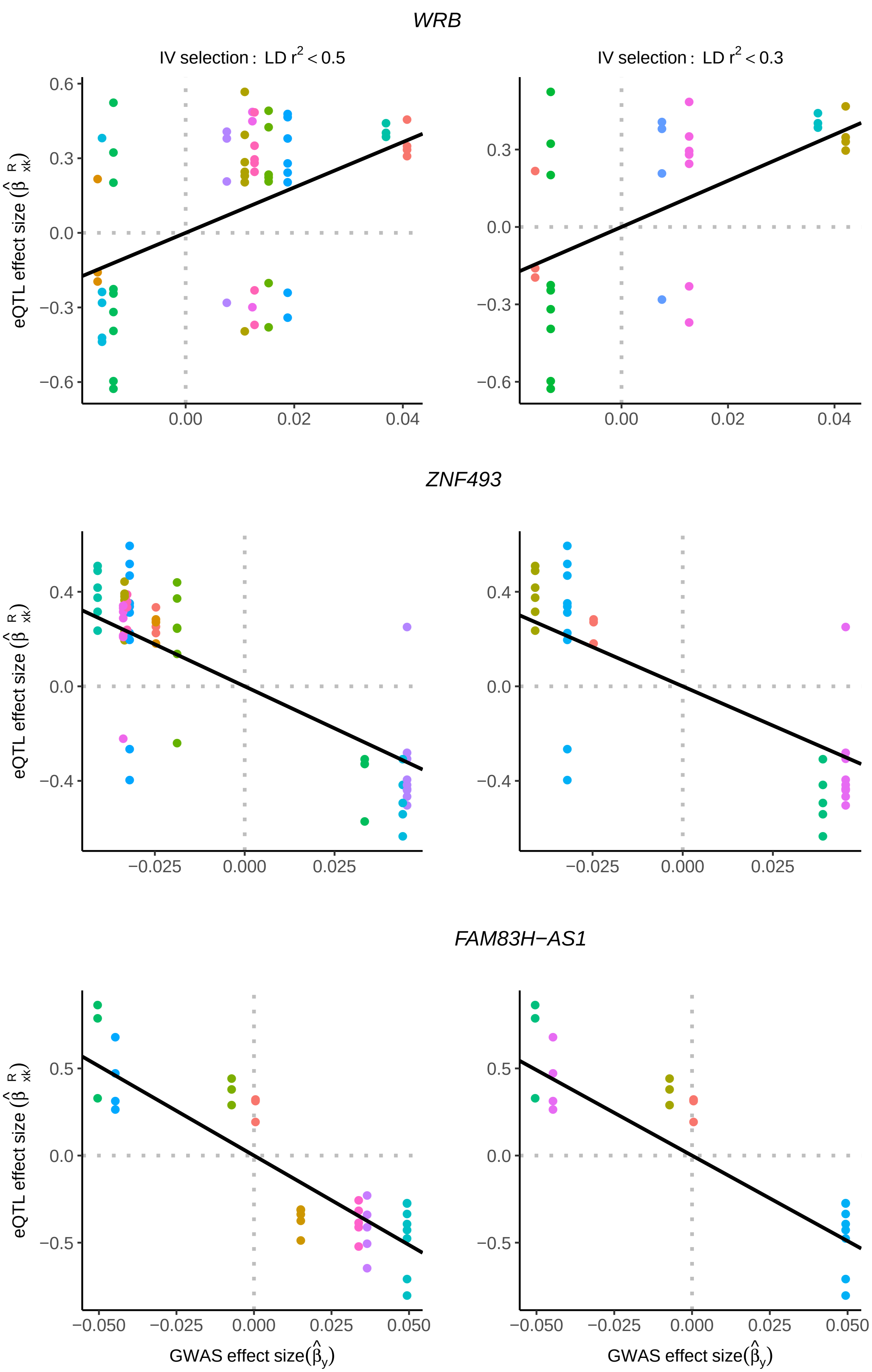
WRB
IV selection : LD r² < 0.5
IV selection : LD r² < 0.3
ZNF493
FAM83H−AS1
eQTL effect size
GWAS effect size

1

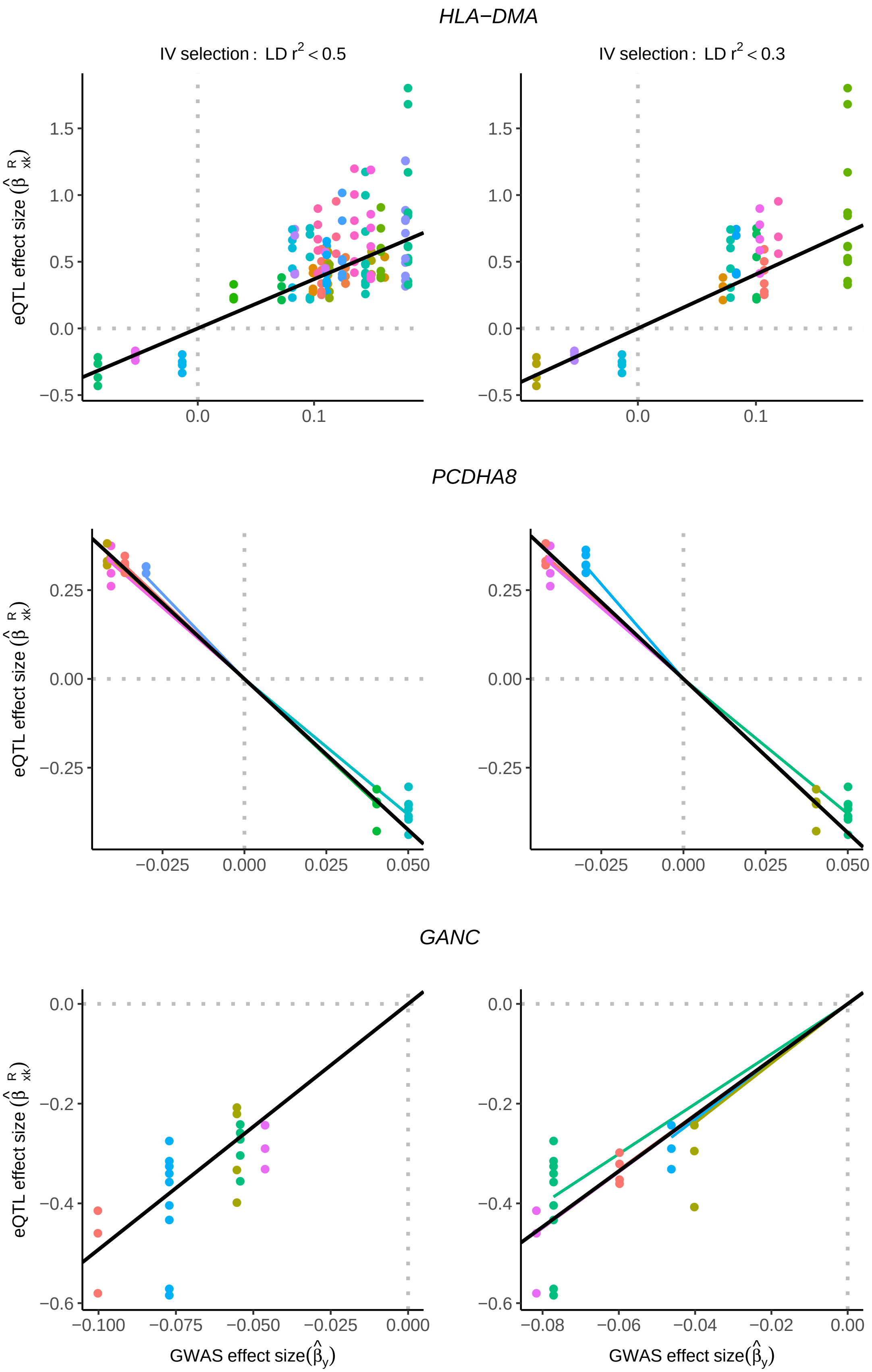
HLA−DMA
IV selection : LD r2 < 0.5
IV selection : LD r2 < 0.3
PCDHA8
GANC
eQTL effect size
GWAS effect size
1.5
1.0
0.5
0.0
−0.5
0.1
0.25
0.00
−0.25
−0.025
0.000
0.025
0.050
−0.2
−0.4
−0.6
−0.100
−0.075
−0.050
−0.08
−0.06
−0.04
−0.02
0.00

1

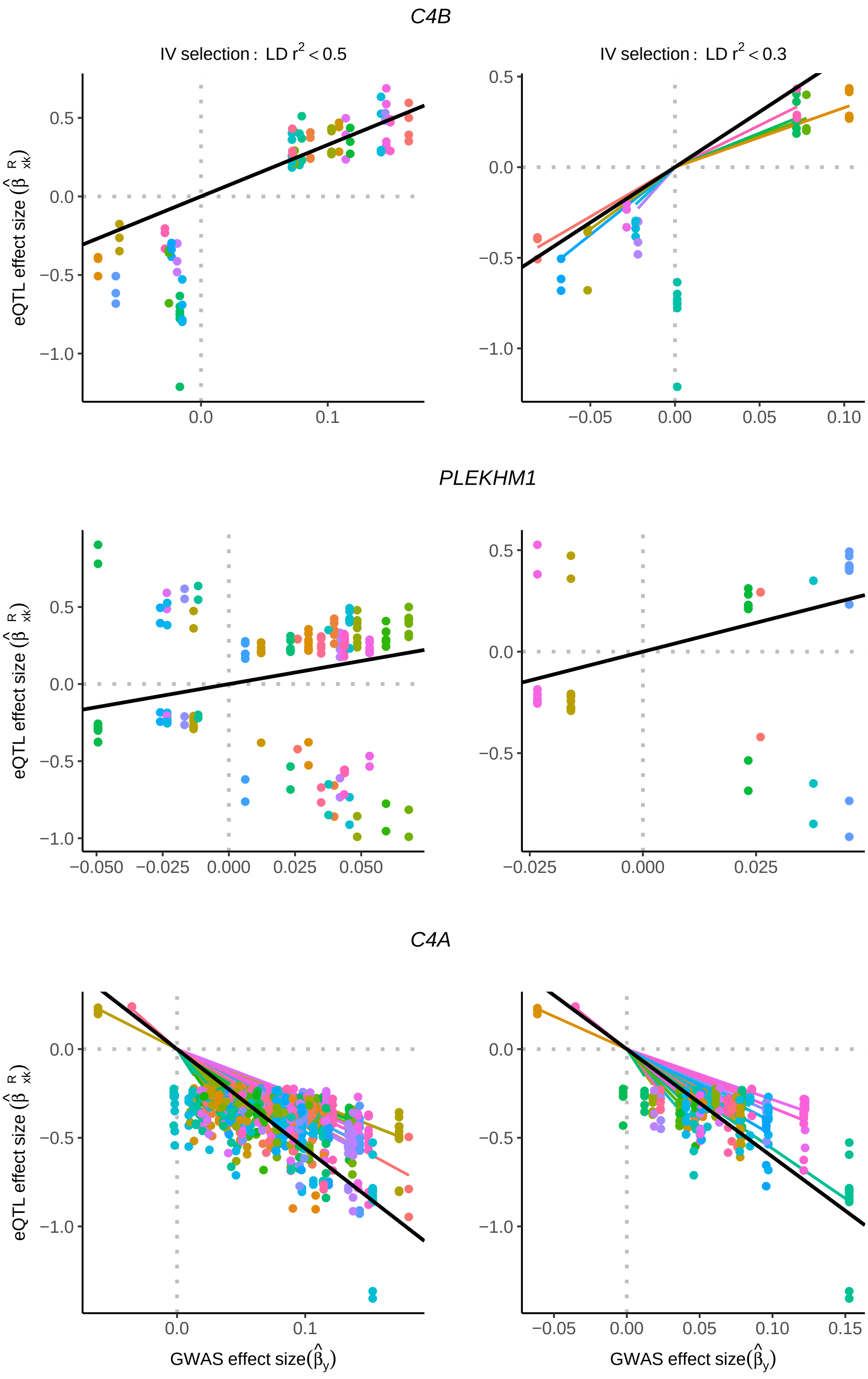

C4B
IV selection : LD r^2 < 0.5
IV selection : LD r^2 < 0.3
PLEKHM1
C4A
eQTL effect size
GWAS effect size
